# Ultrasonic reporters of calcium for deep tissue imaging of cellular signals

**DOI:** 10.1101/2023.11.09.566364

**Authors:** Zhiyang Jin, Anupama Lakshmanan, Ruby Zhang, Teresa A. Tran, Claire Rabut, Przemysław Dutka, Mengtong Duan, Robert C. Hurt, Dina Malounda, Yuxing Yao, Mikhail G. Shapiro

**Author notes:** Corresponding authors: MGS. These authors contributed equally (Anupama Lakshmanan, Ruby Zhang).

## Abstract

Calcium imaging has enabled major biological discoveries. However, the scattering of light by tissue limits the use of standard fluorescent calcium indicators in living animals. To address this limitation, we introduce the first genetically encoded ultrasonic reporter of calcium (URoC). Based on a unique class of air-filled protein nanostructures called gas vesicles, we engineered URoC to produce elevated nonlinear ultrasound signal upon binding to calcium ions. With URoC expressed in mammalian cells, we demonstrate noninvasive ultrasound imaging of calcium signaling *in vivo* during drug-induced receptor activation. URoC brings the depth and resolution advantages of ultrasound to the *in vivo* imaging of dynamic cellular function and paves the way for acoustic biosensing of a broader variety of biological signals.

## INTRODUCTION

Calcium is a ubiquitous signaling molecule involved in essential cellular functions in various tissues and organisms, including synaptic transmission in neurons^1,2^, insulin regulation in islet cells^3,4^, immune activation in T-cells^5–7^ and cytotoxicity in tumors^8,9^. Due to the importance of this signaling molecule, calcium imaging with state-of-the-art synthetic and genetically encoded calcium indicators (GECIs) has proven to be of immense value in biological research^7,10–15^. However, light scattering in tissue makes it challenging to use fluorescent calcium sensors at scale in intact organisms^16^. In most cases, it is limited to single-cell imaging in small, optically clear, or surgically accessed tissues, or lower-sensitivity and lower-resolution imaging in larger and deeper areas in non-transparent organisms^17^. The ability to image calcium at sufficient resolution and depth in the context of intact living organisms would promote our understanding of fundamental physiology and facilitate the development of novel diagnostic and therapeutic agents.

Compared to other prevalent non-invasive imaging modalities such as magnetic resonance imaging^18,19^ and photoacoustics^15,20,21^, ultrasound provides an unparalleled combination of penetration depth (centimeters), imaging volume (multiple cm^3^), spatiotemporal resolution (∼100 µm and ∼1 ms^22,23^), accessibility, and compatibility with freely moving experimental subjects^23^. Recently, the first genetically encoded ultrasound contrast agents were introduced based on gas vesicles (GVs), a unique class of air-filled protein nanostructures derived from buoyant microbes^24^. GVs produce ultrasound contrast due to the low density and high compressibility of their gaseous core relative to aqueous tissues^24^, and their heterologous expression in bacteria^25,26^ or mammalian cells^26,27^ allows GVs to serve as acoustic reporter genes (ARGs).

GVs’ ultrasound contrast depends on the composition and mechanics of their protein shell^28,29^. For example, an alpha-helical protein called GvpC sits on the outside of the shell and stiffens the GV against deformation under acoustic pressure; removal of this protein results in reversible GV buckling and enhanced nonlinear contrast ^28–32^. The first GV-based acoustic biosensors – of protease activity – were developed by engineering GvpC to contain specific protease recognition sites, such that cleavage by the cognate protease makes the GV shell more flexible, leading to stronger nonlinear contrast^33^. While these protease sensors herald an important advance in biomolecular ultrasound^22,23,34^, they produce only a one-time, irreversible change in acoustic contrast due to the permanent covalent modification of the sensor and were only shown to function in bacteria.

Here, we set out to develop the first dynamic, reversible, allosteric acoustic biosensor – an ultrasonic reporter of calcium (URoC) – to enable noninvasive, deep-tissue calcium imaging in mammalian cells (**Fig. 1a**). Inspired by fluorescent GECIs^35^, we hypothesized that we could engineer GvpC to incorporate calmodulin (CaM) and a calmodulin-binding peptide (CBP), such that calcium binding would result in a reversible conformational change that weakens GvpC binding to the GV shell. This would make the GV more flexible and increase its nonlinear acoustic response (**Fig. 1a-b)**. After engineering and systematically characterizing a range of URoC designs, we obtained an acoustic biosensor that produces a 4.7-fold increase in nonlinear contrast in response to calcium *in vitro*, with a sensitivity midpoint of 113 nM. We showed that this construct can be fully genetically encoded and functional in mammalian cells, producing a contrast enhancement of more than 170% in response to elevated intracellular calcium. To validate the performance of the URoC *in vivo*, we engineer cells to increase intracellular calcium through drug-induced activation of a G protein-coupled receptor (GPCR) and imaged the pharmacodynamic response of these cells deep within the mouse brain. This calcium dynamics could be visualized through the intact skull with spatiotemporal resolution of 100 µm and ∼0.6 s. This first-generation URoC demonstrates that acoustic biosensors can be engineered to respond reversibly to intracellular signals, opening the window for ultrasound to follow dynamic cellular processes within the native context of intact living animals.

**Figure 1.**
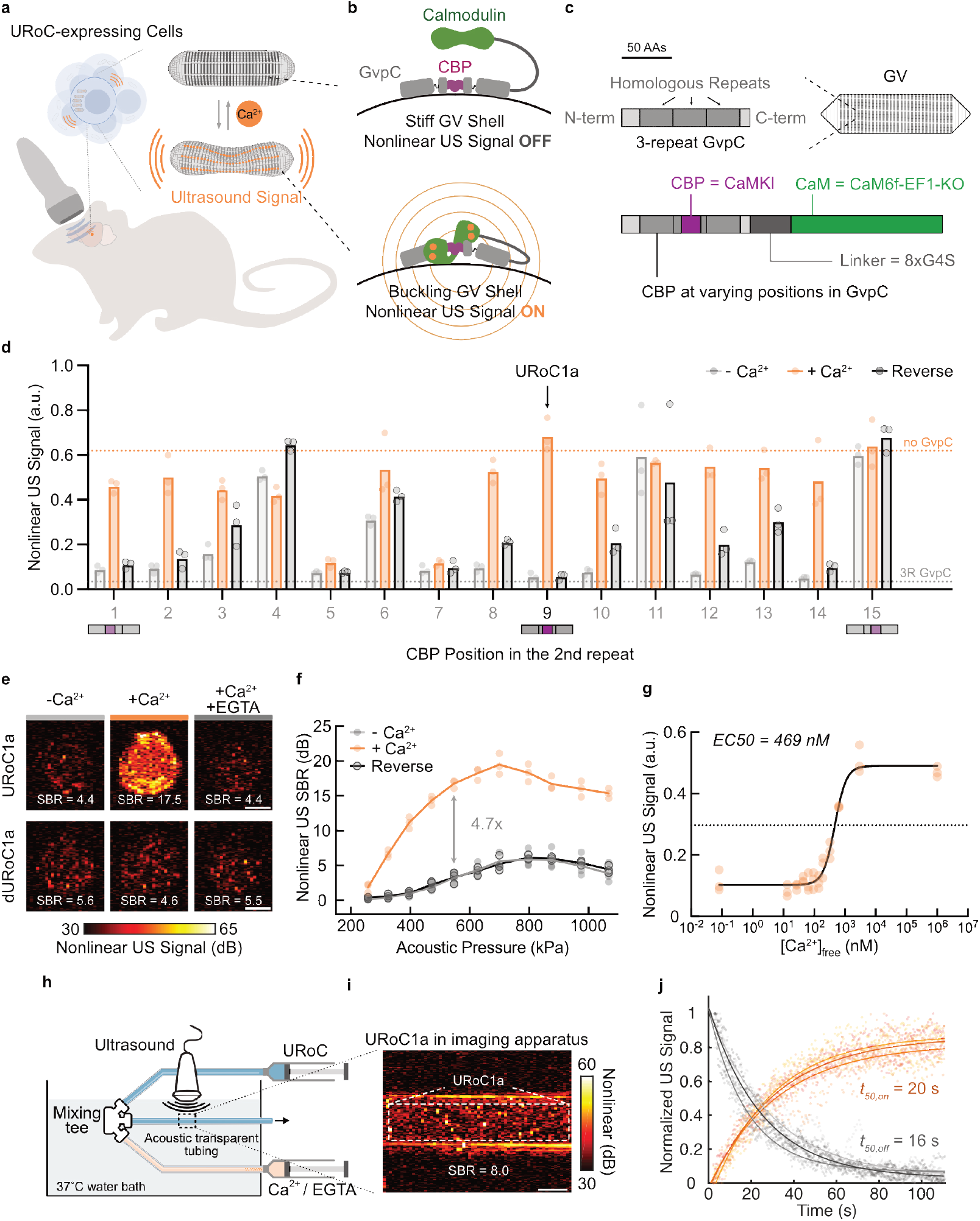
Ultrasonic reporter of calcium dynamics. (**a**) Schematics of URoC transducing intracellular calcium dynamics to ultrasound signal, enabling deep-tissue imaging of cellular function with ultrasound. (**b**) Schematic of URoC and its proposed mechanism of action. (**c**) Top: schematic of a 3-repeat Ana GvpC structure, comprising three 33-amino acid repeats flanked by N- and C-terminal regions. Bottom: schematic of the molecular design of URoC and its engineering components. (**d**) Nonlinear signal after background subtraction and GV concentration normalization of URoC variants with different CBP insertion positions in the second repeat. Samples at OD_500_ 1.8 were incubated with 200 µM CaCl_2_, 5 mM EGTA, or first with 200 µM CaCl_2_ and then with 5 mM EGTA at 37°C before imaging with xAM at 547 kPa. The dash lines indicate the mean signal of GVs with 3R GvpC (gray) or without GvpC (orange). (**e**) Representative ultrasound images of agarose phantom containing URoC1a or control dURoC1a GVs at OD_500_ 1.8 with 200 µM CaCl_2_, 5 mM EGTA, or first with 200 µM CaCl_2_ and then with 5 mM EGTA. The images were acquired with xAM at 547 kPa. (**f**) Nonlinear signal-to-background ratio (SBR) in dB scale as a function of applied acoustic pressure for URoC1a after incubation with 200 µM CaCl_2_, 5 mM EGTA, or first with 200 µM CaCl_2,_ and then with 5 mM EGTA. Solid curves represent the mean of all biological replicates. (**g**) Nonlinear signal of URoC1a as a function of free calcium concentration. The legend lists the EC50 of URoC1a determined from fitting a 4-variable Hill equation. Solid line represents the fitted curve. (**h**) Schematics of the stopped-flow imaging apparatus for measuring URoC kinetics. **(i**) Representative ultrasound image of calcium-saturated URoC1a in acoustically transparent imaging tubing in the apparatus, acquired with xAM at 472 kPa. (**j**) Min-max normalized nonlinear ultrasound signal of URoC1a as a function of time after being mixed into 1 mM CaCl_2_ or 5 mM EGTA. Data from each biological replicates were normalized and fitted to an exponential function and the legend lists the mean half-rise and half-decay time calculated from the fitting. Solid lines represent fitted curves from each biological replicate. Dots represent individual measurements of each replicate for **d, g**, and **j**. Dots represent the mean of two technical replicates for **f**. Scale bars = 1 mm. Color bars represent nonlinear ultrasound signal intensity in the dB scale. N = 3 biological replicates for (**d-g, j**) and each N consists of 2 technical replicates for (**e**-**f**) and 3 for (**j**).

## RESULTS

### Design and characterization of URoC

As the first step, we designed the GvpC to undergo a calcium-dependent conformational change that alters its binding to the GV shell and, consequently, its ability to strengthen the GV. Inspired by the fluorescent GECI^35^, our URoC design comprises a calmodulin-binding peptide (CBP) incorporated in the middle of GvpC’s alpha-helical structure and calmodulin (CaM) at its C-terminus (**Fig. 1b**). We hypothesized that the calcium-free CaM would have negligible effect on GvpC-GV binding, while Ca^2+^-bound CaM would interact with the CBP, triggering the engineered GvpC to undergo an allosteric conformational change resulting in reduced GV shell stiffness, increased buckling, and elevated non-linear ultrasound contrast **(Fig. 1b**). This mechanism is expected to be fully reversible and compatible with continuous imaging of calcium dynamics.

Based on previous experience with GvpC engineering^33^, we developed our URoC GvpC by replacing certain segments of the second repeat of a 3-repeat Ana GvpC (3R GvpC) (**Fig. 1c**) and fused CaM to the C-terminus of the GvpC using a flexible linker (**Fig. 1b-c**). We built and tested a library of 15 initial URoC designs, incorporating the CBP from CaMKI^36^ at different positions within the second GvpC repeat and a mutant CaM from GCaMP6f^11,37^ (CaM6f-EF1-KO) fused to the C-terminus of GvpC using a long flexible linker (8 repeats of G4S, or GGGGS) (**Fig. 1c**). We first tested these GvpC designs using purified GVs isolated from *Anabaena flos-aquae* (Ana) by biochemically replacing their GvpC^38^. We tested the Ca^2+^-dependent nonlinear ultrasound contrast of these engineered GV sensors at 37°C using a cross-propagating amplitude modulation pulse sequence (xAM)^30^. Prior to imaging, samples were incubated for 10 minutes at 37°C with 200 µM Ca^2+^, 5 mM of the calcium chelator EGTA, or first with 200 µM Ca^2+^ and then reversed with 5 mM EGTA. As controls, GVs with unmodified 3R GvpC or no GvpC were tested under the same conditions and showed virtually no calcium-dependent nonlinear contrast (**Supplementary Fig. 1a**).

Most of our 15 variants strengthened the shell in the calcium-free condition, showing low nonlinear signal similar to control GVs with 3R GvpC (**Fig. 1d, Supplementary Fig. 1a**). Nine variants (#1-3, #8-10, #12-14) showed significant enhancement in nonlinear contrast in response to calcium (**Fig. 1d, Supplementary Fig. 1b**). However, among these nine, only variants #1 and #9 showed a complete return to the calcium-free baseline upon sequestration of free Ca^2+^ by EGTA (**Fig. 1d, Supplementary Fig. 1c**), demonstrating fully reversibility (within 10 minutes, see Methods). Of these two, the construct with the CBP at position #9 showed the largest dynamic range – its nonlinear signal in calcium-free or EGTA reversal condition was as low as the GV controls with 3R GvpC, while its signal in the calcium-saturated condition was similar to GVs lacking GvpC (**Fig. 1d, Supplementary Fig. 1a**). We selected this construct for further characterization, naming it URoC1a.

Consistent with results from the initial screening, URoC1a showed reversible ultrasound contrast in response to calcium at the physiological temperature of 37° C, with an enhancement of approximately 13.5 decibels (dB), or a fold change of 4.7x, in nonlinear ultrasound signal-to-background ratio (SBR) (**Fig. 1e-f**). A control “dead” variant of URoC1a – dURoC1a – with all its calcium-binding EF-hands disabled through point mutations, produced low nonlinear signals without any Ca^2+^-dependence (**Fig. 1e Supplementary Fig. 1d**). URoC1a showed minimal sensitivity to magnesium (**Supplementary Fig. 1e)**, while maintaining full calcium-sensing functionality within the physiological range of intracellular pH between 7.0 and 7.5^39^ (**Supplementary Fig. 1f**), with diminished performance outside this range, similar to fluorescent GECIs^35,40^.

We characterized URoC1a sensitivity to calcium and measured a half-maximal response (EC50) at around 469 nM (**Fig. 1g**), within the physiological range of calcium concentration inside mammalian cells^41^. To measure the kinetics of URoC1a, we built a custom stopped-flow ultrasound imaging apparatus where the URoC1a solution and calcium or EGTA solution were equilibrated to 37° C, then delivered into a mixing tee connector, rapidly mixed, and injected into acoustically transparent tubing for imaging with ultrasound (**Fig. 1h-i**). With this apparatus, we were able to measure the kinetics of URoC with a dead time down to 2 seconds (see Methods, **Supplementary Fig. 1g**), and found that the calcium-saturated half-rise time of URoC1a is ∼ 16 seconds, while the half-decay time for reversal with EGTA is around 20 seconds (**Fig. 1j**). These results established URoC1a as an acoustic biosensor for calcium with a large dynamic range, physiologically relevant calcium sensitivity, and kinetics on the order of 10 seconds, with potential for further optimization.

### Mechanism and optimization of URoC

To improve the performance of URoC, especially its kinetics, we first examined its molecular mechanism of action. In our initial hypothesis, the engineered GvpC would partially dissociate from the GV shell upon calcium binding while part of GvpC would act as an anchor to keep it in the proximity of the shell. However, we discovered that the calcium-bound URoC GvpC fully dissociated from the shell, as shown by both cryo-electron microscopy (cryo-EM) and gel electrophoresis (**Fig. 2a, Supplementary Fig. 2a-b**). In cryo-EM images of the longitudinal cross-section of the URoC1a GVs, we observed a dot-shape GvpC density above the α2 helices of GvpA in samples incubated without calcium before freezing (**Fig. 2a**). In contrast, when the URoC1a GVs were pre-incubated with calcium, the density of the GvpC disappeared (**Fig. 2a**), representing a dissociation of GvpC from the GV shell. This finding was confirmed by denaturing gel electrophoresis, which also showed the disappearance of GvpC from buoyancy-purified (see Methods) GVs after calcium incubation (**Fig. 2a**). The control dURoC1a GVs, on the other hand, showed no signs of GvpC dissociation in the presence and absence of calcium, as seen in both cryo-EM and gel electrophoresis experiments (**Supplementary Fig. 2a-b**).

**Figure 2.**
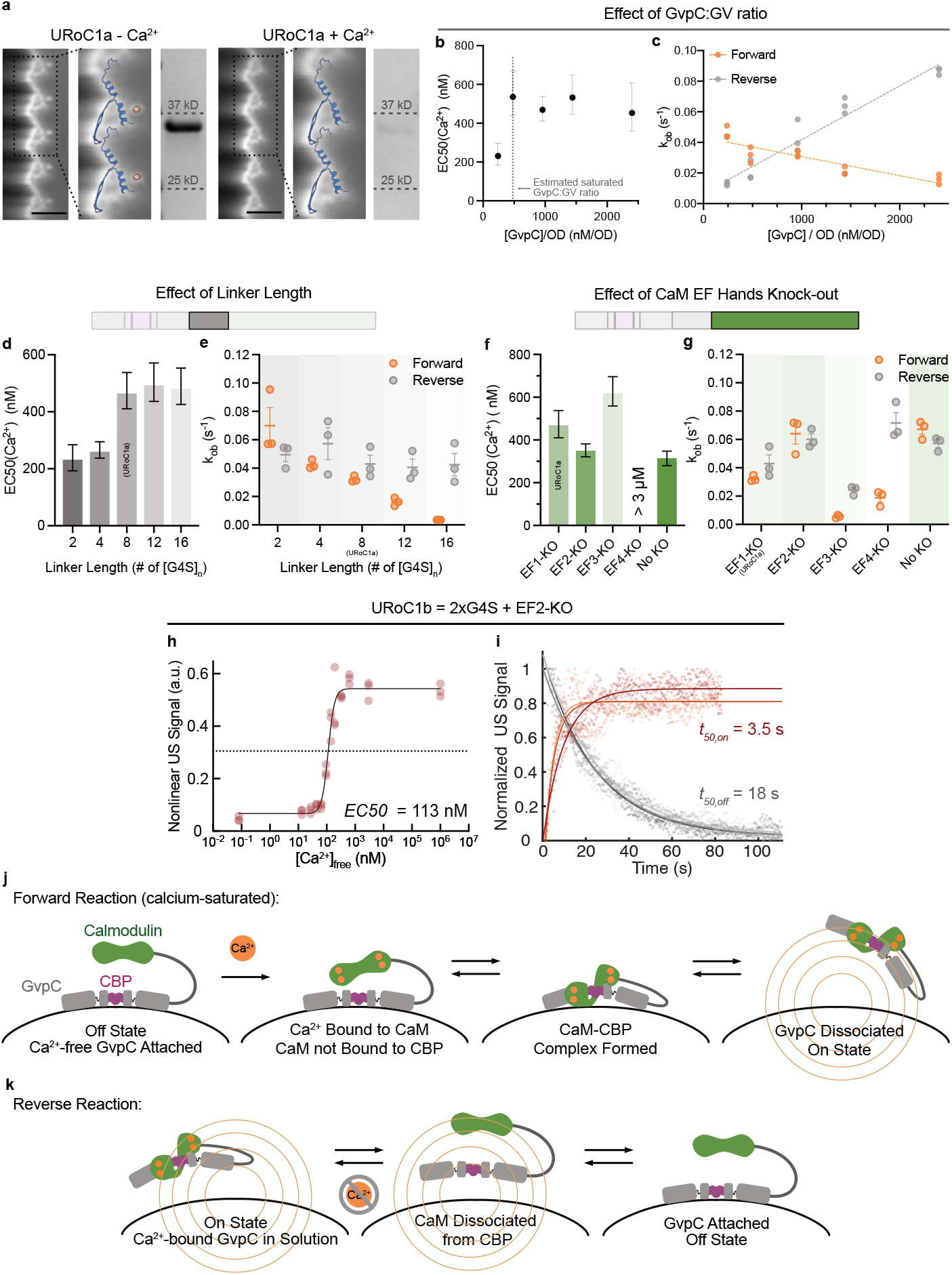
Molecular mechanisms and optimization of URoC. (**a**) Left of each column: cryo-EM 2D density map of the side view of the URoC1a GV shell incubated with 200 µM CaCl_2_ or 5 mM EGTA prior to freezing and an integrative model of the Ana GvpA:GvpC (PDB: 8GBS) complex was overlaid on the GV shell density in the 2D class averages. Right of each column: Coomassie-stained SDS-PAGE gel of OD_500nm_-matched URoC1a GVs with unbound GvpC molecules removed through buoyancy purification after incubation with calcium or EGTA at 37°C. The band corresponds to the size of the URoC1a GvpC. (**b**) The EC50 of URoC1a as a ratio of GvpC concentration to GV concentration (OD_500_). The dashed line indicates the estimated GvpC concentration needed to saturate all the docking sites on GVs. (**c**) The observed rate constants of URoC1a as a function of GvpC concentration relative to GV concentration (OD_500_), for the forward (orange) and reverse (gray) reactions. The data were fitted to a linear function and the dashed lines represent the fitted curves. (**d-e**) The EC50(**d**) and observed rate constants(**e**) of URoC variants as a function of the length of the flexible linker tethering the C-terminus of GvpC to the N-terminus of CaM. (**f-g**) The EC50(**f**) and observed rate constants(**g**) of URoC variants as a function of individual EF-hand knock-outs of the CaM. For the EF4-KO in (**f**), it did not reach its half-maximal contrast at the second highest calcium concentration (3 µM) used in our calcium titration experiments. (**h**) Nonlinear signal of URoC1b as a function of free calcium concentration. N = 3 biological replicates. The legend lists the EC50 of URoC1b determined from fitting a 4-variable Hill equation. Dots represent individual measurements of each replicate and solid line represents the fitted curve. (**i**) Min-max normalized nonlinear ultrasound signal of URoC1b as a function of time after being mixed with 1 mM CaCl_2_ or 5 mM EGTA. N = 3 biological replicates, each with 3 technical replicates. Data from each of the biological replicates were normalized and fitted to an exponential function. The legend lists the mean half-rise and half-decay time calculated from the fitting. Dots represent individual measurements of each replicate and solid lines represent fitted curves from each biological replicate. (**j, k**) Schematics of the proposed molecular mechanism of URoC for the calcium-saturated forward reaction (**j**) and reverse reaction from the calcium-saturated state to the no calcium state (**k**). Ultrasound data was acquired at 547 kPa for (**b, d, f, h**) and at 472 kPa for (**c, e, g, i**). Scale bars = 5 nm for **a**. Dots represent fitted rate constants from each of the biological replicates for **c, e**, and **g**. Error bars represent 95% confidence interval (CI) estimated from the fitting for **b, d, f**, and standard error of the mean (SEM) for **e** and **g**.

After determining that calcium-dependent nonlinear ultrasound contrast originates from the complete dissociation of GvpC, we postulated that the sensitivity and kinetics of URoC1a should depend on the concentration of GvpC relative to its available docking sites on GVs. Indeed, we found that the calcium EC50 of URoC1a decreased to 231 nM with a GvpC concentration of 0.5x relative to the estimated docking sites^28^ (**Fig. 2b**). In comparison, once the GvpC concentration exceeded the docking site concentration, the EC50 stayed almost constant around 469 nM (**Fig. 2b**). In kinetics measurements, increasing the relative GvpC concentration resulted in lower rate constants for the forward reaction (GvpC dissociation), while accelerating the reverse reaction with a larger slope (**Fig. 2c**). In both cases, the rate constant showed reasonably linear dependence on GvpC concentration (R squared = 0.77 and 0.95 for the forward and reverse reaction, respectively). These results pointed to calcium-dependent GvpC dissociation and reassociation being rate-limiting steps in sensor function.

Building on these insights, we tested several modifications in the molecular design of URoC1. We started with the length of the linker connecting CaM to GvpC, which presumably dictates the local concentration of CaM near the CBP. Characterizing URoC variants with different lengths of flexible G4S linkers, we found that shorter linkers resulted in higher calcium sensitivity (**Fig. 2d**). A clear step-change was observed between 4xG4S and 8xG4S, where the EC50 for the variants with 2xG4S or 4xG4S were 236 nM and 264 nM respectively, while those for variants with longer linkers were all slightly below 500 nM. Using an estimated length of 3.8 Å per amino acid for G4S^42^, this step-change happened between 76 Å and 152 Å. Fittingly, we estimated the distance between the C-terminus of the URoC GvpC and the N-terminus of the CaM in the calcium-bound state to be around 92 Å, assuming GvpC holds a rigid alpha-helical structure (see Methods) (**Supplementary Fig. 2c**). We hypothesize that when the linker is shorter than this length, the calcium-bound URoC GvpC may adopt a conformation where part of the helical structure of GvpC is disrupted leading to lower affinity to the GV shell and thus higher sensitivity to calcium. We also found that shorter linkers provided faster forward kinetics, where the shortest 2xG4S linker improved the sensor kinetics by approximately a factor of 2 to a half-rise time at 10 seconds (**Fig. 2e**), while linker length had a negligible effect on reverse kinetics.

Additionally, we tested how EF-hand mutations on the CaM affect the URoC performance, as such mutations were shown to improve the kinetics of certain fluorescent GECIs^37^. We constructed URoC variants with each of the four EF-hands disabled through point mutations (EF*x*-KO, *x* = 1-4) and with the intact CaM from GCaMP6f^11^ (No-KO). The EF2-KO and No-KO variants showed similar characteristics, both with better sensitivity and faster kinetics than the URoC1a with EF1-KO (**Fig. 2f-g**). The EC50 for EF2-KO and No-KO were 350 nM and 315 nM, respectively, and the half-rise time of both variants decreased to ∼10 seconds, with the reverse kinetics insignificantly faster than that of URoC1a (**Fig. 2f-g**). The EF3-KO and EF4-KO variants had lower sensitivity and slower forward kinetics. These results suggest that the second half of CaM (C lobe) may be more dominant than the first half (N lobe) in the actuation of the URoC GvpC.

Hypothesizing that the effects of the linker length and CaM mutations can be synergetic, we combined the shortest linker (2xG4S) and the CaM with the second EF-hand knocked out (EF2-KO) to create URoC1b. As predicted, both the sensitivity and forward kinetics of URoC1b were substantially improved, with an EC50 of 113 nM and a half-rise time of just 3.5 seconds (**Fig. 2h-i**), approaching the limits of our stopped-flow kinetics measurement. In the meantime, there was no significant change in the reverse rate constant (half-decay ∼18 seconds) (**Fig. 2i**). These results provide a more sensitive and faster biosensor – URoC1b – and showcase how the modular design of URoCs may enable the future construction of acoustic biosensors with different calcium sensitivity and kinetics for various applications, following a path similar to optical GECIs^43^.

Our optimization experiments lead us to a conceptual model of URoC operation to facilitate future engineering. For the forward “on” reaction in response to calcium, the binding of calcium-activated CaM and CBP could be rate-limiting, since we observed that the effectively higher local concentration of CaM in constructs with shorter linkers accelerated the sensor’s response (**Fig. 2e,j**). In addition, the production of nonlinear contrast may be rate-limited by GvpC dissociation from the GV shell, evidenced by the inverse relationship between excess free GvpC concentration and forward response rates (**Fig. 2c, j**). Meanwhile, for the reverse reaction after the removal of free calcium, we identified the re-association of GvpC to the GV shell as a major rate-limiting factor, with baseline return accelerated with higher GvpC concentration (**Fig. 2c, k**). In addition, since some EF-hands mutations in the CaM (without altering the GvpC segment) led to dramatically slower reverse kinetics, we speculate that CBP dissociation from the CaM could also be rate-limiting (**Fig. 2g, k**). We consider calcium-binding to have a minor contribution to the URoC’s response time, since CaM alone binds to calcium ions and changes its conformation within tens of milliseconds^44,45^ and ever faster with the CBP of CaMKI^46^. Our optimization results, combined with this hypothetical reaction scheme, provide a basis for future URoC engineering.

### Ultrasound imaging of calcium dynamics in mammalian cells

After demonstrating the performance of URoCs *in vitro* using purified proteins, we endeavored to demonstrate their functionality inside mammalian cells. Recently, genetic constructs were developed for robust mammalian expression of Ana GVs, enabling ultrasound imaging of gene expression^26^. To express URoCs in mammalian cells as intracellular sensors, we generated Ana GV constructs co-expressing the URoC GvpC along with the other genes encoding Ana GVs. These constructs include one polycistronic plasmid encoding the assembly factor genes *gvpNJKFGWV* linked through P2A self-cleaving peptides and a second plasmid encoding the main structural protein GvpA, followed by an internal ribosome entry site (IRES)^47^ and the URoC GvpC (**Fig. 3a**). This architecture allowed us to maintain a constant GvpA to GvpC ratio while tuning their ratio relative to assembly factors for robust expression^26^.

**Figure 3.**
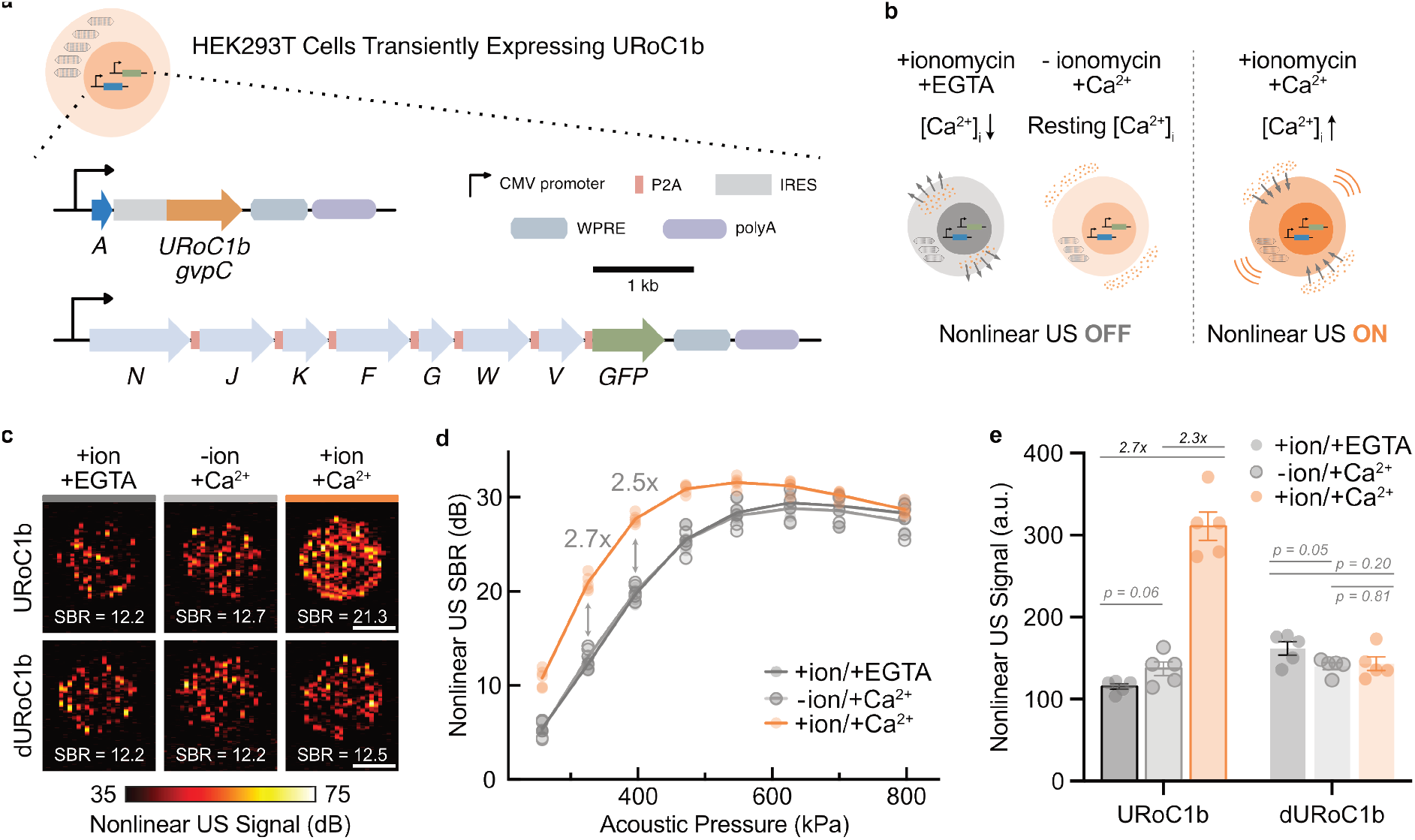
Ultrasound imaging of calcium in mammalian cells. (**a**) Schematics of the genetic constructs for transient expression of URoC1b in HEK293T cells. (**b**) Schematics of applying URoC1b to image ionomycin-induced intracellular calcium signaling. (**c**) Representative ultrasound images of agarose phantom containing cells expressing URoC1b or control dURoC1b GVs at 5 million cells per mL with ionomycin and EGTA, ionomycin, and calcium or calcium only. (**d**) Nonlinear SBR in dB scale as a function of applied acoustic pressure for cells expressing URoC1b after incubation with ionomycin and EGTA, ionomycin, and calcium or calcium only. (**e**) Nonlinear signal of cells expressing URoC1b in phantoms with ionomycin and EGTA, ionomycin and calcium, or calcium only. Ultrasound data were acquired with xAM at 326 kPa for (**c, e**). Dots represent the mean of two technical replicates for **d, e**. N = 5 biological replicates with each N consisting of 2 technical replicates for **d** and **e**. Solid curves represent the mean of all replicates. The bars indicate the mean of all replicates. Scale bars = 1 mm. Color bars represent nonlinear ultrasound signal intensity in the dB scale.

We expressed URoC1b in HEK293T cells, an established cell line for biosensor development^10^. We transiently transfected the cells with our two plasmids, induced changes in their intracellular calcium concentration [Ca^2+^]_i_ with the ionophore ionomycin^48,49^ and imaged them with nonlinear ultrasound. We expected [Ca^2+^]_i_ elevation to above 1 µM upon incubation with ionomycin^49^ and calcium to induce maximal nonlinear signal from the calcium-saturated URoC, and [Ca^2+^]_i_ depletion with ionomycin and EGTA to induce minimal signal (**Fig. 3b**). We also tested cells incubated with physiological calcium without ionomycin to test sensor response to the resting state [Ca^2+^]_i_ (**Fig. 3b**).

As hypothesized, cells incubated with ionomycin and calcium showed higher nonlinear ultrasound contrast across a range of acoustic pressures compared to cells incubated with ionomycin and EGTA, with the peak dynamic range at 326 kPa showing a SBR enhancement of 9 dB, or a fold change of 2.7x in nonlinear signal (**Fig. 3c-e**). Relative to cells with physiological resting state, the [Ca^2+^]_i_ ionomycin incubation provided an 8 dB increase or a 2.3x fold change in SBR (**Fig. 3c-e**). As additional controls, we evaluated cells transfected with the mutated calcium-insensitive sensor dURoC1b, which did not produce any significant [Ca^2+^]_i_-dependent contrast (**Fig. 3c-e, Supplementary Fig. 3a**).

To evaluate the tolerability of URoC1b expression, we measured the viability of cells expressing the biosensor and found no substantial difference compared to cells expressing the fluorescent GECI jRCaMP1b^50^ (**Supplementary Fig. 3b**). Taken together, these results demonstrate the first ultrasound imaging of calcium dynamics in mammalian cells and present URoC1b as a robust tool for cellular calcium imaging.

### Ultrasound imaging of drug-induced calcium signaling in vivo

After successfully establishing the basic design principles of URoC and characterizing their performance *in vitro* and inside the cells, we assessed its capacity for imaging physiologically relevant calcium dynamics *in vivo*. As a model system, we chose GPCR-driven calcium signaling, which plays critical functional roles in many cell types^51–54^. In particular, G_q_-coupled GPCRs elevate [Ca^2+^]_i_ through inositol triphosphate (IP_3_) signaling to the endoplasmic reticulum (ER), leading to the cytoplasmic release of ER calcium^54,55^. To demonstrate *in vivo* URoC performance in a tightly controlled model system, we generated a stable HEK293T cell line co-expressing URoC1b with the hM3D(G_q_) DREADD, a muscarinic receptor engineered to respond to bio-orthogonal ligands such as descloroclozapine (DCZ)^56,57^. We chose HEK293T cells for this purpose due to their relatively low endogenous GPCR signaling^58^ and previous use as *in vivo* cell-based biosensors^58–60^. We hypothesized that by implanting these cells in tissue such as in the brain, we would be able to follow the calcium response of these cells to a systemically administered ligand such as DCZ, thereby monitoring its pharmacodynamics (**Fig. 4a**). In addition to providing a well-controlled proof of concept for *in vivo* URoC functionality, such monitoring has intrinsic utility for drug and receptor engineering.

**Figure 4.**
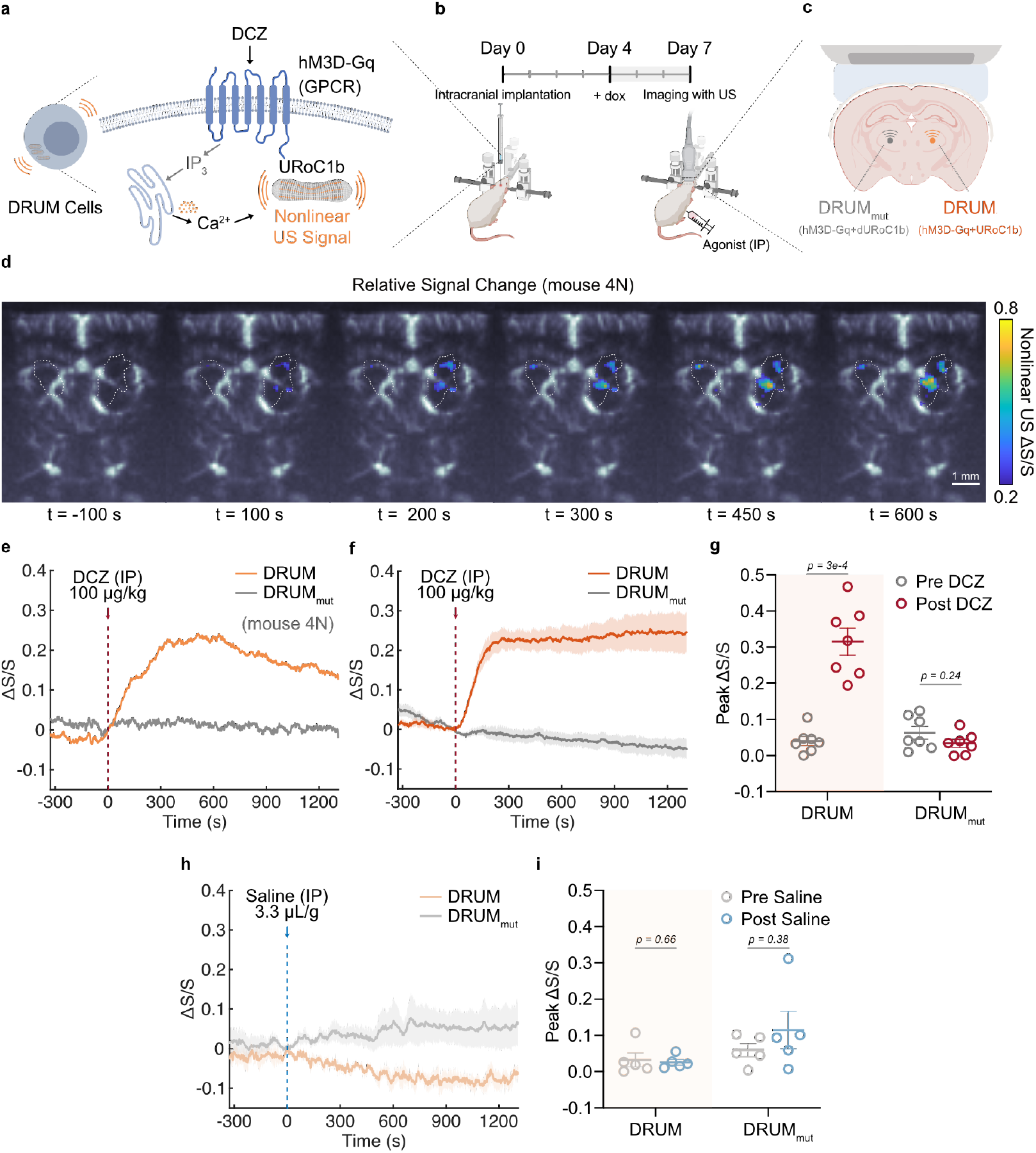
Ultrasound imaging of ligand-induced GPCR-driven calcium dynamics in vivo. (**a**) Schematics of engineered cells stably expressing hM3D(G_q_) and URoC1b as DRUM cells. (**b**) Schematics of timeline for intracranial implantation, induction for URoC1b expression, and ultrasound imaging. (**c**) Schematics of the implant layout of DRUM and DRUM_mut_ cells in the mouse brain. (**d**) Representative nonlinear US images of DRUM and DRUM_mut_ implants in the brain of mouse 4N. The relative signal change at t = - 100/100/200/300/450/600 s compared to t = 0 s was overlaid on a Doppler image. The white dash lines represent the contour of the regions that showed nonlinear signal above the noise threshold (see Methods) at t = 0. The Scale bar = 1 mm. Color bars represent ultrasound signal intensity in the dB scale. The intensity of the Doppler image represents the blood volume that circulated in each voxel within an integration time of 400 ms (see Methods). (**e**-**f, h**) Time traces of nonlinear ultrasound contrast of the implanted cells. DCZ (**e**-**f**) or saline (**h**) was injected intraperitoneally (I.P.) at t = 0 s at the dashed line. The nonlinear ultrasound signal was acquired with xAM at 426 kPa (calibrated in water). For (**e**), Individual time trace for the same mouse 4N in (**d**) was shown. (**g, i**) Peak relative signal change of DRUM and DRUM_mut_ implants before and after DCZ (**g**) or saline (**i**) stimulation. For (**f, h**), dark curve = mean and shaded region = SEM. For (**g, i**), lines = mean and error bars = SEM. N = 7 mice for the DCZ stimulation (**f, g**) and N = 5 mice for the saline control (**h, i**). Imaging was conducted through polymer cranial windows for (**d**-**i)**.

We constructed a stable HEK293T cell line expressing the hM3D(G_q_) receptor through lentiviral infection, then used the PiggyBac transposase system^61^ to add doxycycline-inducible^62^ genes encoding URoC1b (**Fig. 4a**), resulting in the DRUM (Drug-Receptor Ultrasound Monitoring) cells. In parallel, we used the non-calcium sensing mutant dURoC1b to produce the DRUM_mut_ cells. After generating the cell lines, we implanted DRUM cells unilaterally into the mouse thalamus, with DRUM_mut_ cells implanted contralaterally. (**Fig. 4b-c**). Seven days after implantation and 3 days after starting daily doxycycline induction, we performed nonlinear ultrasound imaging through an acoustically transparent polymer cranial window^63,64^ (**Fig. 4b**). We located the DRUM and DRUM_mut_ cells through their baseline nonlinear contrast, imaged their baseline signal for 5 minutes, then administered intraperitoneal DCZ (100 µg/kg) and continued to acquire ultrasound images over 20 minutes (**Fig. 4b**). At the end of the imaging session, we acquired a Doppler vasculature map as an anatomic reference.

The nonlinear signal of the DRUM cells started to increase shortly after systemic DCZ administration, reaching a plateau around 350 seconds and remaining steady over the remaining 20 min (**Fig. 4d-e, Supplementary Video 1**), consistent with DCZ pharmacodynamics^57^. Our images easily spatially resolved the DRUM and contralateral DRUM_mut_ cells. Within the DRUM implant, we observed region-specific kinetics, where certain locations showed a decreasing signal trend after reaching the peak while others remained steady after reaching a plateau (**Fig. 4d, Supplementary Fig. 4a-c**), potentially due to differences in vascularization, receptor expression, downstream signaling, and desensitization^65^. In addition, there were parts of the DRUM implant that did not show any response to the stimulation, potentially due to inhomogeneity in GPCR expression or vascular accessibility.

Immunofluorescence imaging of brain tissue collected after the imaging session showed good alignment of ultrasound signal and fluorescence from the implanted cells (**Supplementary Fig. 5a-b**), confirming expression of both the URoCs and the hM3D(G_q_), while also revealing variability in hM3D(G_q_) expression and some necrotic areas around in which the ligand might not be easily accessible (**Supplementary Fig. 5a**), consistent with our ultrasound imaging data.

Across 7 mice, DRUM cells demonstrated a significant increase of approximately 25% in nonlinear ultrasound signal (**Fig. 4f-g, Supplementary Fig. 6a**) upon the administration of DCZ, with the average baseline fluctuation of 1% (**Supplementary Fig. 6b**). Our sensors detected a characteristic onset time of around 128 seconds (**Fig. 4f, Supplementary Fig. 6a**), similar to the 3 minutes previously estimated with invasive measurements^57^. The DRUM_mut_ implants did not show a substantial response to DCZ, and both the DRUM and DRUM_mut_ implants showed no significant response in control saline injections (**Fig. 4h-i, Supplementary Fig. 6c**). With a similar DCZ stimulation experiment, we also tested the performance of DRUM implants transcranially and showed in 5 mice that we could achieve a 22% peak enhancement with similar kinetics through the intact skull without any additional surgeries (**Supplementary Fig. 6d-e**). These results demonstrate the ability of URoC1b to noninvasively visualize the spatiotemporal dynamics of drug-induced, receptor-mediated calcium signaling across a large area deep in the mouse brain, thus validating the basic *in vivo* ultrasound imaging capabilities of this technology.

## DISCUSSION

Taken together, our results establish a paradigm for visualizing intracellular calcium dynamics with ultrasound. This paradigm is enabled by engineering the GV-stiffening protein GvpC to undergo a calcium-dependent, allosteric conformational change, leading to increased nonlinear ultrasound signal. To materialize this concept, we engineered and optimized a series of URoCs, resulting in a biosensor with large dynamic range, high sensitivity, full reversibility, and functionality inside mammalian cells.

The *in vivo* application demonstrated in this work provides a fundamental proof of concept for real-time ultrasound calcium imaging inside a living animal. As an immediate application, the DRUM approach could be used to map the region-specific pharmacodynamics of various chemogenetic ligands and correlate it with behavioral responses or neuroimaging (e.g. with functional ultrasound^64,66,67^). Similar cellular constructs could be developed to image the action of a large variety of pharmaceutical agents or drugs of abuse.

Additional work is needed to adapt URoCs to a wider range of calcium imaging applications. With parallel, ongoing advances in gene delivery methods and GV gene cluster engineering, it should be possible to express URoCs in primary cells such as neurons *in vivo* in the future. This would enable brain-wide calcium imaging in genetically defined neuronal populations in intact animals, providing an unprecedented view of brain circuit function. Likewise, expression of URoCs in pancreatic beta cells or immune cells would enable the *in vivo* imaging of critical aspects of endocrine and immune activity in basic biology and facilitate the development and monitoring of cell-based therapies. With these and other potential applications, URoCs could have a transformative impact on diverse areas of biology and medicine.

Following the initial invention of calcium biosensors based on fluorescent proteins, GECIs were optimized over multiple generations to achieve their current outstanding performance, and we anticipate a similar development path for URoCs. Although the first-generation biosensors presented in this study have sufficient performance for certain applications, there is ample room for improvement, most notably in their kinetics. Inspired by the approaches taken to improve GECIs, future work could optimize URoCs through rational engineering and directed evolution of components such as the CBP-CaM pair, fusion positions and linkers, and the underlying GV proteins GvpC and GvpA. For example, based on our mechanistic understanding, engineering the calcium-bound GvpC to remain partly anchored to the GVs is expected to improve kinetics. Further improvements could arise from engineering the stoichiometry and identity of GvpC, GvpA and other GV genes^68^, leading to URoC GVs with varying size, shape or mechanical properties. In parallel with molecular engineering, improvements in ultrasound instruments and pulse sequences, such as 3D volumetric imaging^67^, wearable long-term probes ^69,70^, advanced nonlinear imaging and super-resolution^71^ will provide new and improved ways to visualize URoCs *in vivo*. This will mirror the co-evolution of GECIs with advances in optical microscopy^72–74^.

Going beyond calcium imaging, we anticipate that URoCs will serve as a template for the development of other acoustic biosensors based on allosteric conformational changes in GvpC. making it possible to sense a larger variety of ions, enzymes, and other biological signals, helping ultrasound rock and roll toward new breakthroughs in biology and medicine.

## MATERIAL AND METHODS

### Design and cloning of genetic constructs

All the plasmids in this study were constructed through a combination of polymerase chain reaction (PCR) using Q5 polymerase and Gibson assembly^75^ or KLD mutagenesis. All the reagents were from New England Biolabs (NEB) and custom primers were from Integrated DNA Technologies (IDT). For screening and characterization of purified proteins, all *gvpC* gene sequences were codon-optimized for *E. coli* expression and driven by a T7 promoter and lac operator in a pET28a expression vector (Novagen). Specifically, the construct encoding the 3-repeat WT Ana *gvpC* gene was first generated through deletion of the fourth and fifth repeats from an existing plasmid encoding the 5-repeat WT Ana *gvpC* gene with a C-terminus 6xHis tag through KLD mutagenesis. Second, the codon-optimized CaM sequence from GCaMP6f (synthesized by IDT) and an 8xG4S linker sequence were introduced through Gibson assembly. Last, the sequence of CBP from CaMKI was introduced by substitution/insertion into the second repeat of the *gvpC* gene sequence through KLD mutagenesis to generate the URoC *gvpC* genes. Additional modifications such as changing the linker length or CaM mutations were also achieved through KLD mutagenesis.

For the transient expression of URoC in mammalian cells, all sequences were codon-optimized for expression in human cells and driven by a CMV promoter in a pCMVSport vector with WPRE-hGH polyA. Similar steps were taken to modify the WT *gvpC* sequence to the URoC1b/control *gvpC*, which were then introduced along with an IRES sequence into an existing plasmid encoding the *gvpA* gene (Addgene #197588) through Gibson assembly. An existing plasmid was used for the expression of all 7 chaperones (Addgene #197589). For generating the DRUM or DRUM_mut_ stable cell lines, the lentiviral transfer plasmid constitutively expressing hM3D(G_q_)-mCherry was constructed as follows: the sequence encoding hM3D(Gq)-mCherry was amplified by PCR from a plasmid from Addgene (#50474) and assembled into a lentiviral backbone with a human EF1α promoter, an IRES followed by a puromycin resistance gene (*PuroR*) for selection and a WPRE element. The PiggyBac transposon plasmids for URoC expression were constructed by PCR-amplifying the region between the start codon of *gvpNJKFGWV* or *gvpA*-IRES-*gvpC* and the end of the hGH polyA from the transient plasmids. The amplified regions were assembled into the PiggyBac transposon backbone (System Biosciences) between a TRE3G promoter (Takara Bio) for doxycycline-inducible expression and a constitutive EF1α core promoter driving a blasticidin resistance gene *BSD* or hygromycin resistance gene *HygR* gene for selection. The complete list and source of plasmids used in this study are given in **Supplementary Table 1**. All the plasmids were cloned using NEB Turbo *E. coli* (New England Biolabs) and sequence verified.

### Preparation of purified GVs for *in vitro* assays

For the *in vitro* purified protein assays, GVs were collected and purified from confluent Ana cultures using previously published protocols^28,38^. The concentration of Ana GVs was determined by measurement of their optical density (OD) at 500 nm (OD500) using a Nanodrop spectrophotometer (Thermo Fisher Scientific), using the resuspension buffer as the blank. As established in previous work^76^, the concentration of GVs at OD500 = 1 is approximately 184 pM and the gas fraction is 0.0417%. The engineered GvpC was expressed and purified following a previously published protocol with minor modifications. Briefly, for the expression and purification of URoC GvpC variants, plasmids were transformed into chemically competent BL21(DE3) cells (Invitrogen) and grown overnight for 12–16 h at 37°C in 5 ml starter cultures in 2xYT medium with 50 μg/mL of kanamycin. Starter cultures were inoculated 1:100 into auto-induction Terrific Broth (Novagen 71491) with 50 μg/mL of kanamycin and allowed to grow at 30°C (250 r.p.m. shaking) for 20-24 hours for protein expression. Cells were then collected by centrifugation at 5,500 RCF and lysed at room temperature using SoluLyse (Amsbio L200125), Supplemented with protease inhibitor cocktail (10 µL per 1 mL of culture, Sigma P8849), lysozyme (400 μg/mL) and DNase I (10 μg/mL). GvpC inclusion bodies were isolated by centrifugation at 15,000 RCF for 10 min and then resuspended in a solubilization buffer comprising 20 mM of Tris-HCl buffer with 500 mM of NaCl and 6 M of urea (pH: 8.0), before his-tag purification with Ni-NTA (QIAGEN) and wash and elution buffers of the same composition as the solubilization buffer, but with 20 mM and 250 mM imidazole, respectively. The concentration of the purified protein was assayed using the Bradford Reagent (Bio-rad).

Engineered GVs with the URoC GvpC or wild-type GvpC were prepared using published protocols with urea stripping and GvpC re-addition^28,38^. Briefly, Ana GVs were stripped of their native GvpC through treatment with a 6 M urea solution buffered with 100 mM Tris-HCl (pH:8-8.5), followed by two rounds of centrifugally assisted floatation with removal of the subnatant liquid after each round. The recombinant engineered GvpC variants purified from inclusion bodies were then mixed with the stripped Ana GVs in 6 M urea with varying molar excess concentration determined after accounting for 1:15 binding ratio of 3-repeat GvpC:GvpA. For an *n*-fold (2-fold unless specified) stoichiometric excess of GvpC relative to binding sites on an average Ana GV, the concentration of recombinant GvpC (in nmol) to be added to stripped GVs was calculated according to the formula: *n* * OD_500nm_ * 480 nM. The mixture of stripped GVs (OD_500nm_ = 4) and recombinant GvpC in 6 M urea buffer was loaded into dialysis pouches made of regenerated cellulose membrane with a 6-8 kDa M.W. cutoff (Spectrum Labs). The GvpC was allowed to slowly refold onto the surface of the stripped GVs by dialysis in 4 L of calcium-free MOPS buffer (30 mM MOPS, 100 mM KCl, pH 7.2) with 1 mM EDTA for the first round, followed by a second round of dialysis in 4 L of MOPS buffer with 30 µM EGTA. Both rounds were done at 4°C for 8-12 hours each. The dialyzed GV samples were directly used for further characterization.

### *In vitro* assay for calcium response

All data were acquired at 37°C unless specified. For the steady-state calcium response experiments including pH and magnesium sensitivity, engineered GVs at OD_500_ = 3.6 were mixed 1:1 with 1% w/v agarose (Lonza, #50070) in MOPS buffer with 400 µM CaCl_2_ or 10 mM EGTA (final OD_500_ = 1.8) and then loaded into hydrogel phantoms made of 1% agarose in MOPS buffer with 200 µM CaCl_2_ or 5 mM EGTA, followed by incubation and ultrasound imaging at 37°C. For the calcium-free and calcium-saturated conditions, GV samples were loaded directly into the phantoms for incubation. For the reversal condition, 1 M CaCl_2_ solution was first added to the GV samples after dialysis to make a final concentration of 200 µM CaCl_2_. The samples then were incubated at 37°C for 10 minutes before being 1:1 mixed with 1% agarose in MOPS buffer with 10 mM EGTA and loaded into a phantom containing 5 mM EGTA. After loading, the phantoms with GV samples were incubated at 37°C for 10 minutes and then imaged by ultrasound.

For the calcium titration experiments, the zero free calcium buffer (30 mM MOPS, 100 mM KCl, 10 mM EGTA, pH 7.2) and 39 µM free calcium buffer (30 mM MOPS, 100 mM KCl, 10 mM CaEGTA, pH 7.2) were made in lab and serial dilution was performed to generate buffers with different free calcium concentrations. These buffers were calibrated, with and without 1:1 dilution with MOPS with 30uM EGTA, using fluo-4 (Invitrogen F14200) and a commercial calcium calibration buffer (Invitrogen C3008MP). GV samples at final OD_500_ = 1.8 were loaded into the phantoms made of 1% agarose (w/v, Lonza, #50070) in those buffers and incubated at 37°C for >10 minutes before ultrasound imaging.

For measurements of kinetics, GV samples at OD_500_ = 3.6 and the MOPS buffer with 2 mM CaCl_2_ (forward reaction) or 10 mM EGTA (reverse reaction) were loaded into 1 mL syringes that were both controlled by a syringe pump (Kent Scientific). For the reverse reaction, samples were first incubated with 200 µM [Ca^2+^] for > 30 minutes at 37°C before the measurement. The samples and buffer were first delivered into a silicone tubing (OD 1/8”, ID 1/16”) for prewarming at 37°C for 2 minutes. Next, both solutions were injected into a 1:1 mixing tip (MIXPAC T-mixer, Medmix) at a flow rate of 100 µL/s (post-mixing at 200 µL/s) and then into a 1/12” polyolefin tubing for ultrasound imaging after mixing. All the tubing was immersed in a water bath at 37°C. The URoC1a samples pre-incubated with 1 mM CaCl_2_ were mixed with MOPS buffer with 1 mM CaCl_2_ to estimate the dead time. The practical dead time was estimated to be around 2 seconds, considering the following two effects. Dead volume of the setup (150 uL) generates dead time of 0.75 s; Additionally, it took approximately 1.2 seconds for the flow to fully stop (**Supplementary Figure. 1d**).

### Cryo-EM characterization and image analysis

A sample of Ana GVs with addition of GvpC variants was incubated for 30 min at 37°C in presence of 200 µM CaCl_2_ or 5 mM EGTA. A 3 μL volume of sample was applied to C-Flat 2/2 −3C grids (Protochips) that were glow-discharged (Pelco EasiGlow, 10 mA, 1 min). GV samples were frozen using a Mark IV Vitrobot (FEI, now Thermo Fisher Scientific) (37°C, 60% humidity, blot force 3, blot time 4 s). Before applying the GV sample, the grid was incubated in the Vitrobot chamber for 5 minutes to equilibrate to 37°C. Movie stacks were acquired on a 300 kV Titan Krios microscope (Thermo Fisher Scientific) equipped with a K3 6k × 4k direct electron detector (Gatan). Multi-frame images were collected using SerialEM 3.39 software^77^. Approximately 20 movies were acquired for each condition at a pixel size of 1.4 Å (64,000× magnification) with varying defocus from - 1.0 to - 3.0 μm. To generate side-view averages of the GV shell, acquired movies were imported into cryoSPARC^78^. Motion and CTF corrected micrographs were used for subsequent data processing. Using template matching, a few thousand particles of the GV shell’s edges were picked, extracted, and subjected to iterative 2D classification. For visualization purposes, an integrative model of the Ana GvpA:GvpC (PDB: 8GBS)^31^ complex was overlaid on the GV shell density in the 2D class averages.

### Denaturing polyacrylamide gel electrophoresis (SDS-PAGE)

GV samples were incubated with 200 µM CaCl_2_ or 5 mM EGTA for 30 minutes at 37°C and then centrifuged at 300 RCF for 2-4 hours at 37°C. The subnatant liquid was aspirated to remove any unbound GvpC molecules, and the floating layer of GVs were resuspended in MOPS buffer with 200 µM CaCl_2_ or 5 mM EGTA. This step was repeated for 3 times and the samples were concentration-matched at OD_500nm_ = 10 and mixed 1:1 with 2x Laemmli buffer (Bio-Rad), containing SDS and 2-mercaptoethanol. The samples were then boiled at 95°C for 5 minutes and 20 μL of the samples were loaded into a pre-made polyacrylamide gel (Bio-Rad) immersed in 1x Tris-Glycine-SDS Buffer. 10 uL of Precision Plus Protein™ Dual Color Standards (Bio-Rad) was loaded as the ladder. Electrophoresis was performed at 120V for 55 minutes, after which the gel was washed in DI water for 15 minutes to remove excess SDS and coomassie-stained for 1 hour on a rocker using the SimplyBlue SafeStain (Invitrogen). The gel was allowed to de-stain overnight in DI water before imaging using a Bio-Rad ChemiDoc™ imaging system.

### Structure prediction of URoC GvpC

The structure of the GvpC segment of URoC1a (1-127) was predicted using AlphaFold2 (ColabFold^79^) and the structure with the highest confidence was used to align with the existing structure of the CaMKI-CaM complex (PDB: 1MXE)^36^. After the alignment, the distance was measured between the second carbon atom of the G127 in the GvpC and the first carbon atom of the L1 of the CaM using ChimeraX^80^.

### HEK293T cell culture and transient transfection

HEK293T cells (American Type Culture Collection (ATCC), CLR-2316) were seeded in 6-well plates at 2.5 × 10^5^ cells per well at 37 °C and 5% CO_2_ in a humidified incubator in 2 ml of DMEM (Corning, 10-013-CV) with 10% FBS (Gibco), 10 mM HEPES (Cytia) and 1× penicillin–streptomycin 24 hours before transfection. Transient transfection mixtures were created by mixing 2 µg of plasmid mixture with polyethyleneimine (PEI-MAX, Polysciences) at 4 μg of polyethyleneimine per microgram of DNA. The mixture was incubated for 12 minutes at room temperature and added drop-wise to HEK293T cells. Media was changed after 12–16 hours and daily thereafter. For the transient expression of URoC1b and dURoC1b, control GVs, 280 fmol of the *gvpA*-IRES-*gvpC* plasmid and 70 fmol of the plasmid encoding *gvpNJKFGWV* was added into the DNA mixture, and pUC19 plasmid DNA was supplemented to make the total amount of DNA 2 µg. For the transient expression of jRCaMP1b, DNA of the same mass as the GV genes (i.e., *gvpA*-IRES-*gvpC* and *gvpNJKFGWV*) was mixed with the pUC19 plasmid DNA to make the total mass 2 µg. Transfected cells were assayed 72 hours after the transfection.

### Preparation of cells for ionomycin stimulation

After 3 days of expression, cells were dissociated using Trypsin/EDTA (Corning 25-053-CI), centrifuged at 300 RCF for 6 minutes at room temperature and resuspended in HEPES-buffered Hanks’ balanced salt solution without calcium (HHBSS, 20 mM HEPES, 2 g/L D-glucose in 1x HBSS without calcium). The cells were counted using an automated cell counter (Countess™ 3, Thermo Fisher) and all the samples were concentration-matched to 10 million cells per milliliter in HHBSS without calcium. HHBSS phantoms were prepared with 1% agarose (w/v, Lonza, #50070) supplemented with 10 µM ionomycin (Sigma, I9657) and 5 mM EGTA, 10 µM ionomycin and 2 mM CaCl2, or 2 mM CaCl2. Cells were diluted 1:1 with the 1% agarose containing 2x concentration of the corresponding reagents to result in a final concentration of 5 million cells per milliliter before loading into their respective phantoms. The phantoms were incubated at 37°C for 10 minutes before ultrasound imaging in a water bath with the same buffer content at 37°C.

### *In vitro* ultrasound imaging

As described above, GV or cell samples were loaded into imaging phantoms made of 1% agarose (w/v) in specified buffers or injected into an acoustically transparent tubing for imaging (**Fig. 1h**). All the samples were placed in the same buffer as the phantom during the imaging session and maintained at 37°C by a custom water bath unless specified.

Imaging was performed using a Verasonics Vantage programmable ultrasound scanning system and a L22-14vX 128-element linear array Verasonics transducer, with a specified pitch of 0.1 mm, an elevation focus of 8 mm, an elevation aperture of 1.5mm and a center frequency of 18.5 MHz with 67% −6 dB bandwidth. For nonlinear image acquisition, a custom cross-amplitude modulation (xAM) sequence detailed in an earlier study^30^ was used with an xAM angle (θ) of 19.5°, an aperture of 65 elements, and a transmitting frequency at 15.625 MHz. The center of the sample wells was placed at a depth of 5 mm with a conventional ray-line scanning B-mode pulse sequence with parabolic focusing at 10 mm and an aperture of 40 elements. The focus was set to be far from the sample position to reduce the acoustic pressure to avoid collapsing the samples. The transmitted pressure at the sample position at 5 mm was calibrated using a Precision Acoustics fiber-optic hydrophone system. Each image was an average of 50 accumulations. B-mode images were acquired at a transmit voltage of 1.6V (86 kPa), and an automated voltage ramp imaging script (programmed in MATLAB) was used to conduct xAM acquisitions at different acoustic pressures. For purified GV samples, an xAM voltage ramp sequence from 3V (258 kPa) to 8V (1068 kPa) in 0.5V step increments was used. For HEK293T cells expressing GVs, an xAM voltage ramp sequence from 3 V (258 kPa) to 6.5V (797 kPa) in 0.5V step increments was used. Samples were subjected to complete collapse at 25V with the B-mode sequence for 10 seconds, and the subsequent post-collapse xAM images acquired at the same voltage steps were used as the blank for data processing. For purified GV samples, B-mode images at the beginning of the pressure ramp and at the end of the post-collapse ramp were acquired for concentration normalization.

### Viability assay

The viability of the cells transfected with URoC1b and jRCaMP1b was assayed with alamarBlue™ (Invitrogen DAL1025) for resazurin reduction, the luminescent ATP detection assay kit (Abcam, ab113849) for the ATP content and Trypan Blue for the membrane permeability, all following the manufacturers’ protocols. In particular, for the reducing power assay, the cells were incubated with the alamarBlue™ reagent for 1 hour at 37°C and 5% CO_2_ before the read-out with a plate reader (Tecan). For the Trypan Blue assay, cells were first dissociated using Trypsin/EDTA (Corning 25-053-CI), and the culture media with 10% FBS was added to the wells at a 1:1 ratio to quench the reaction. Cells were resuspended in the trypsin-media mixture and then proceeded for viability quantification using an automated cell counter (Countess™ 3, Thermo Fisher) after 1:1 dilution with the Trypan Blue.

### Construction of DRUM and DRUM_mut_ cell lines

HEK293T cells were cultured as described above and seeded in 6-well plates at 2.5 × 10^5^ cells per well. After 24 hours, cells were lentivirally transduced with jmL-95_pEF1α-hM3D(G_q_)-mCherry-IRES-PuroR-WPRE at a multiplicity of infection (MOI) of 4 with 10 µg/mL of polybrene. The media containing the viruses and polybrene was removed 12 hours after the infection and the cells were passaged 36 hours after the infection. Next, the cells were expanded to a surface-treated T-75 flask and treated with 2 µg/mL of puromycin (Invivogen) for 2 weeks for selection of the transduced cells to generate the cell line expressing hM3D(G_q_), HEK-jmL95.

After the puromycin selection, HEK-jmL95 was seeded in 2 separate wells in a 6-well plate at 2.5 × 10^5^ cells per well without puromycin. 22 hours after the seeding, the culture media was replaced with DMEM (Corning, 10-013-CV) containing 2% FBS (Gibco), 10 mM HEPES (Cytia) and 1× penicillin–streptomycin. After 2 hours of incubation with reduced-serum media, cells were transfected with 3 µg of PiggyBac transposon:transposase plasmid mixture (2143 ng of URoC plasmids with a molar ratio of 4:1 PB-gvpA-IRES-URoC1b/dURoC1b:PB-gvpNV transposons and 857 ng of PiggyBac transposase) using PEI-MAX with the same protocol for the transient transfection. Media was changed to 10% FBS culture media after 12-16 hours of incubation and daily thereafter for 3 days. The transfected cells were expanded to T-75 flasks and cultured with 2 µg/mL of puromycin (Invivogen), 10 µg/mL of blasticidin (Invivogen) and 200 µg/mL of hygromycin (Invivogen) for selection of cells transduced with the URoC1b genes. After 2 weeks, the selected cells were expanded in media without puromycin, blasticidin or hygromycin and frozen in Recovery Cell Culture Freezing Medium (Gibco) using Mr. Frosty cell freezing container (Nalgene) filled with isopropanol at −80 °C and then stored in liquid nitrogen (LN2) vapor phase until use.

### Intracranial implantation of DRUM and DRUM_mut_ cells

Transcranial injection of DRUM and DRUM_mut_ cells was conducted in NSG mice (Jackson Laboratory) aged 6 weeks for those imaged transcranially or 6-9 weeks for others imaged through cranial windows, all under a protocol approved by the Institutional Animal Care and Use Committee of the California Institute of Technology. No randomization or blinding was necessary in this study. DRUM and DRUM_mut_ cells were recovered from LN_2_ storage and cultured without drug selection for at least 2 passages before implantation. On the day of implantation, cells were dissociated using Trypsin/EDTA (Corning 25-053-CI), centrifuged at 500 g for 5 minutes, and resuspended in culture media without antibiotics at a concentration of 60 million cells per milliliter. The cells were then stored on ice before injection.

During the surgery, mice were anesthetized with 1–2% isoflurane, weighed before the surgery, maintained at 37 °C on a heating pad, weighed before the surgery, and placed on a stereotaxic instrument. The cells were injected intracranially at the coordinate of Anterior-Posterior (AP) –2 mm, Media-Lateral (ML) ±1.5 mm (–1.5 mm for DRUM_mut_ and +1.5 mm for DRUM cells), and Dorsal-Ventral (DV) –3.5 mm, relative to the bregma. The injection was conducted through a microliter syringe (Hamilton) with a needle of 33G (World Precision Instrument) controlled by a micro syringe pump (World Precision Instrument) at a flow rate of 7 nL per second. A volume of 3300 nL containing 200 thousand cells was injected on each side.

For the mice to be imaged transcranially, the skin was closed after the injection with a tissue adhesive (GLUture). Animals recovered quickly and remained bright, alert, and responsive before the ultrasound imaging.

For the mice to be imaged through cranial windows, the craniotomy was performed after the injection using a micro drill steel burr (Burr number 19007-07, Fine Science Tools) from approximately AP –1 mm to AP –3 mm and from ML –2.5 mm to ML +2.5 mm. A 0.125 mm thick polymethylpentene TPX^®^ film (Sigma) was cut to cover the cranial opening and then attached to the skull through a light-cured composite (Tetric EvoFlow). Dental cement (C&B METABOND) was used to close between the skin and the cranial window.

### Ultrasound imaging of GPCR signaling in vivo

4 days after the implantation and every 12 hours thereafter, the mice were induced for GV expression through intraperitoneally (I.P.) injection of 150 µL of saline containing 150 µg of doxycycline. The ultrasound imaging was conducted 7 days after the surgery (induced for 3 days). The mice were first anesthetized with 4% isoflurane, and then placed on a stereotaxic instrument with 1.5% isoflurane for maintenance anesthesia, and their core temperature was monitored with a rectal probe and maintained at 37°C with a heating pad.

The same instruments and pulse sequences with the *in vitro* ultrasound imaging were used, and the ultrasound transducer L22-14vX was held by a custom holder mounted on the right arm of the stereotaxic instrument.

For the mice with the cranial windows, the ultrasound coupling gel (Aquasonic) was directly applied to the cranial windows, and the transducer surface was placed ∼1 mm above the middle of the cranial window in the coupling gel. The B-mode pulse sequence described in the in vitro imaging methods was applied to fine-tune the initial position of the transducer, and the xAM pulse sequence was applied at 426 kPa (calibrated in water, higher than the optimal pressure in vitro to account for attenuation through the cranial window and tissue) to scan across the cranial window to locate the DRUM and DRUM_mut_ implants via their off-state nonlinear signal. The coronal planes with the highest signal of both implants were chosen for imaging. After locating the imaging plane, the imaging was paused (no ultrasound transmitted or received), with the mice staying in the imaging apparatus for 10 minutes to ensure full equilibrium of mouse body temperature, breathing, and other potential effects of isoflurane.

After 10 minutes, the imaging session started using xAM with the same parameters as described above – 65 elements aperture, 426 kPa (calibrated in water), and a frame rate of approximately 1.7 frames per second. The baseline signal was acquired for 500 frames, after which 100 µg/kg of DCZ (in the format of 30 µg/mL of deschloroclozapine dihydrochloride in saline) or 3.3 µL/g of saline was injected I.P. The nonlinear signal was recorded for 2000 frames after the injection. 7 mice were stimulated with DCZ, and 5 mice were injected with saline as controls. For 4 mice stimulated with DCZ, a vasculature map for anatomic reference was acquired through a plane wave power Doppler pulse sequence at 6V at the end of the imaging session.

For mice imaged transcranially, similar procedures were followed except for three steps. First, the skin was cut open and the ultrasound gel (Aquasonic) was applied to the skull to couple the transducer to the skull. Second, an amplitude modulation pulse sequence with a parabolic focus at 6 mm and an aperture of 40 elements at a higher peak positive pressure of 1.28 MPa (calibrated in water) was used for imaging to account for the attenuation of the intact skull. Third, the plane wave Doppler imaging was acquired at 25V to account for the intact skull. 5 mice were imaged transcranially for the DCZ stimulation.

### Doppler image acquisition

The power Doppler images mapping local changes in cerebral blood volume (CBV) were acquired at 15.625 MHz as previously described^81,82^. Briefly, the pulse sequence contains 15 tilted plane waves varying from −14° to 14° at a 500 Hz pulse repetition frequency. A block of 200 coherently compounded frames was processed using an SVD clutter filter to separate tissue signal from blood signal (cutoff of 40) to obtain a final power Doppler image exhibiting CBV in the whole imaging plane^83,84^.

### Collection of brain tissue

After the imaging session, all the animals were perfused with 30 mL of phosphate-buffered saline (PBS), followed by 30 mL of 10% formalin solution. The brains were resected and placed in 10% formalin solution for 36 hours at 4°C after which they were transferred to PBS for long-term storage at 4°C. The brain tissue for one of the mice stimulated with DCZ (mouse 3N) was accidently lost during sample transportation.

### Histology of implanted cells in mouse brain

A representative brain from a mouse stimulated with DCZ (mouse 4N with the cranial window) was sectioned into 100 µm slices using a vibrating microtome (Leica). The slices were mounted and stained with DAAPI nuclear stain using ProLong Diamond antifade mountant with DAPI (Thermo Fisher P36962) and sealed with acrylic resins. The mounted slices were imaged using a Zeiss LSM 980 confocal microscope with ZEN Blue. Images were processed and exported using the ZEN Blue software.

### Image processing and data analysis

All *in vitro* and *in vivo* ultrasound images were processed using MATLAB. Regions of interest (ROIs) were manually defined so as to adequately capture the signals from each sample well or region of the implants in the brain. For the steady state calcium response, the sample ROI dimensions (1.2 mm × 1.2 mm square) were the same for all *in vitro* phantom experiments except the URoC initial screening and calcium titration experiments, which were centered at each well. The background ROI was manually selected from the background for each pair of sample wells. For the URoC initial screening and calcium titration experiments, 96-well plate layout phantoms with larger well size were used, so ROIs with a size of 2 mm (lateral) × 1 mm (axial) were chosen for analysis. For each ROI, the mean pixel intensity *I*_*ROI*_ was calculated, and the pressure-sensitive ultrasound intensity (Δ*I* = *I*_*intact*_−*I*_*collapsed*_) was calculated by subtracting the mean pixel intensity of the collapsed image from the mean pixel intensity of the intact image. The signal-to-background ratio (SBR) in was calculated for each sample well by 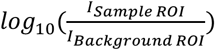. For purified protein characterization, due to the correlation between the GV concentration and B-mode signal^24^, the pressure-sensitive B-mode intensity (Δ*I*_*B* -*mode*_ = *I*_*Bmode,pre-ramp*_− *I*_*Bmode,collapsed*_) was used to normalize the small variability in concentrations. The background-subtracted concentration-normalized nonlinear signal at a specific voltage (or acoustic pressure) was calculated with the following formula: 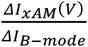. For the kinetics measurement, the ROIs were chosen to cover the interior of the tubing and raw mean intensity was used for min-max normalization within each biological replicates. The data after normalization were fitted to an exponential equation *Ae*^*Bt*^+ *C* with data from the first 3 frame post-mixing not used for curve fitting to account for the dead time (**Supplementary Fig. 1g**). The fitted parameter B was used as the observed rate constant.

For the *in vivo* experiment, rectangle ROIs were selected to contour the brain region with the above-background nonlinear signal, while tissue background ROI was selected at the same depth with the implants. For quantification, the mean intensity within the ROIs were used to calculate the relative signal change as 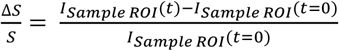 and a moving average filter of size 5 was applied to smooth the curve. For image display, this calculation was done on individual pixels to generate relative change images. Then, a median filter with a [1,4] (lateral, axial) neighborhood and a Gaussian filter with a standard deviation of 1 were applied to reduce the noise of the images. The images were overlaid on top of the Doppler images for display. All images were pseudo-colored (bone colormap for Doppler images, hot colormap for xAM images and parula for the relative signal changes), with the maximum and minimum levels indicated in the accompanying color bars. For the contour showing the location of implants, the threshold was calculated as the mean of the background ROI plus twice its standard deviation based on the image at t = 0 s.

### Statistical analysis

Error bars indicate the mean standard error of the mean (SEM) unless otherwise specified. The sample size is N=3 biological replicates in all *in vitro* experiments unless otherwise stated. For each biological replicate, there were technical replicates to accommodate for variability in experimental procedures such as sample loading and pipetting. SEM was calculated by taking the values for the biological replicates, each of which was the mean of its technical replicates. The numbers of biological and technical replicates were chosen based on preliminary experiments such that they would be sufficient to report significant differences in mean values. P values were calculated using a two-tailed paired t-test.

## Data availability

All plasmids used in this study are available from M.G.S. under a material agreement with the California Institute of Technology. The key genetic constructs will be deposited with Addgene at the time of manuscript publication. Ultrasound acquisition and processing scripts used to generate key figures and results will be posted to a publicly accessible GitHub repository at the time of manuscript publication. Raw data are available from the corresponding author upon reasonable request.

## ACKNOWLEDGEMENTS

The authors thank R. Nayak for providing calibration data for the ultrasound transducer used for imaging. Cryo-Electron microscopy was done in the Beckman Institute Resource Center for Transmission Electron Microscopy at Caltech. The authors also thank X. Chen and V. Gradinaru for sharing instruments and protocols for tissue sectioning. Figure. 4b was created with BioRender.com. This research was supported by the National Institutes of Health (R01NS120828 to M.G.S.) and DARPA (W911NF-14-1-0111). A.L. was supported by the NSF graduate research fellowship (Award No. 1144469) and the Biotechnology Leadership pre-doctoral Training Program in Micro/Nanomedicine (Rosen Bioengineering Center and NIH Training Grant 5T32GM112592-03/04). R.C.H. was supported by the Caltech Center for Environmental Microbial Interactions. Related research in the Shapiro Laboratory is supported by the David and Lucile Packard Foundation and the Chan-Zuckerberg Initiative. M.G.S. is an investigator of the Howard Hughes Medical Institute.

## AUTHOR CONTRIBUTIONS

Z.J., A.L., and M.G.S. conceived and designed the study. Z.J., and A.L. designed and planned experiments. Z.J., A.L., R.Z., T.A.T., C.R., M.D., R.C.H, D.M., and Y.Y. conducted the experiments. Z.J. wrote the scripts for ultrasound imaging and data processing. Z.J. and Y.Y. analyzed the data. Z.J. wrote the manuscript, with input from all authors. M.G.S. supervised the research. All authors have given approval to the final version of the manuscript.

## COMPETING FINANCIAL INTERESTS

Z.J., M.G.S., A.L., and T.A.T. are inventors on one patent application related to this work filed by California Institute of Technology (no. 17/937,975, filed 1 October 2022). The authors declare that they have no other competing financial interests.

## REFERENCES

1. Grienberger, C. & Konnerth, A. Imaging Calcium in Neurons. Neuron 73, 862–885 (2012).

2. Spitzer, N. C., Gu, X. & Olson, E. Action potentials, calcium transients and the control of differentiation of excitable cells. Current Opinion in Neurobiology 4, 70–77 (1994).

3. Rorsman, P., Braun, M. & Zhang, Q. Regulation of calcium in pancreatic α- and β-cells in health and disease. Cell Calcium 51, 300–308 (2012).

4. Campbell, J. E. & Newgard, C. B. Mechanisms controlling pancreatic islet cell function in insulin secretion. Nat Rev Mol Cell Biol 22, 142–158 (2021).

5. Trebak, M. & Kinet, J.-P. Calcium signalling in T cells. Nat Rev Immunol 19, 154–169 (2019).

6. Vig, M. & Kinet, J.-P. Calcium signaling in immune cells. Nat Immunol 10, 21–27 (2009).

7. Thestrup, T., Litzlbauer, J., Bartholomäus, I., Mues, M., Russo, L., Dana, H., Kovalchuk, Y., Liang, Y., Kalamakis, G., Laukat, Y., Becker, S., Witte, G., Geiger, A., Allen, T., Rome, L. C., Chen, T.-W., Kim, D. S., Garaschuk, O., Griesinger, C. & Griesbeck, O. Optimized ratiometric calcium sensors for functional in vivo imaging of neurons and T lymphocytes. Nat Methods 11, 175–182 (2014).

8. Halle, S., Keyser, K. A., Stahl, F. R., Busche, A., Marquardt, A., Zheng, X., Galla, M., Heissmeyer, V., Heller, K., Boelter, J., Wagner, K., Bischoff, Y., Martens, R., Braun, A., Werth, K., Uvarovskii, A., Kempf, H., Meyer-Hermann, M., Arens, R., Kremer, M., Sutter, G., Messerle, M. & Förster, R. In Vivo Killing Capacity of Cytotoxic T Cells Is Limited and Involves Dynamic Interactions and T Cell Cooperativity. Immunity 44, 233–245 (2016).

9. Weigelin, B., den Boer, A. T., Wagena, E., Broen, K., Dolstra, H., de Boer, R. J., Figdor, C. G., Textor, J. & Friedl, P. Cytotoxic T cells are able to efficiently eliminate cancer cells by additive cytotoxicity. Nat Commun 12, 5217 (2021).

10. Tian, L., Hires, S. A., Mao, T., Huber, D., Chiappe, M. E., Chalasani, S. H., Petreanu, L., Akerboom, J., McKinney, S. A., Schreiter, E. R., Bargmann, C. I., Jayaraman, V., Svoboda, K. & Looger, L. L. Imaging neural activity in worms, flies and mice with improved GCaMP calcium indicators. Nat Methods 6, 875–881 (2009).

11. Chen, T.-W., Wardill, T. J., Sun, Y., Pulver, S. R., Renninger, S. L., Baohan, A., Schreiter, E. R., Kerr, R. A., Orger, M. B., Jayaraman, V., Looger, L. L., Svoboda, K. & Kim, D. S. Ultrasensitive fluorescent proteins for imaging neuronal activity. Nature 499, 295–300 (2013).

12. Dana, H., Sun, Y., Mohar, B., Hulse, B. K., Kerlin, A. M., Hasseman, J. P., Tsegaye, G., Tsang, A., Wong, A., Patel, R., Macklin, J. J., Chen, Y., Konnerth, A., Jayaraman, V., Looger, L. L., Schreiter, E. R., Svoboda, K. & Kim, D. S. High-performance calcium sensors for imaging activity in neuronal populations and microcompartments. Nat Methods 16, 649–657 (2019).

13. Zhang, Y., Rózsa, M., Liang, Y., Bushey, D., Wei, Z., Zheng, J., Reep, D., Broussard, G. J., Tsang, A., Tsegaye, G., Narayan, S., Obara, C. J., Lim, J.-X., Patel, R., Zhang, R., Ahrens, M. B., Turner, G. C., Wang, S. S.-H., Korff, W. L.,Schreiter, E. R., Svoboda, K., Hasseman, J. P., Kolb, I. & Looger, L. L. Fast and sensitive GCaMP calcium indicators for imaging neural populations. Nature 615, 884–891 (2023).

14. Su, Y., Walker, J. R., Hall, M. P., Klein, M. A., Wu, X., Encell, L. P., Casey, K. M., Liu, L. X., Hong, G., Lin, M. Z. & Kirkland, T. A. An optimized bioluminescent substrate for non-invasive imaging in the brain. Nat Chem Biol 19, 731–739 (2023).

15. Qian, Y., Piatkevich, K. D., Mc Larney, B., Abdelfattah, A. S., Mehta, S., Murdock, M. H., Gottschalk, S., Molina, R. S., Zhang, W., Chen, Y., Wu, J., Drobizhev, M., Hughes, T. E., Zhang, J., Schreiter, E. R., Shoham, S., Razansky, D., Boyden, E. S. & Campbell, R. E. A genetically encoded near-infrared fluorescent calcium ion indicator. Nat Methods 16, 171–174 (2019).

16. Taroni, P., Pifferi, A., Torricelli, A., Comelli, D. & Cubeddu, R. In vivo absorption and scattering spectroscopy of biological tissues. Photochemical & Photobiological Sciences 2, 124–129 (2003).

17. Girven, K. S. & Sparta, D. R. Probing Deep Brain Circuitry: New Advances in in Vivo Calcium Measurement Strategies. ACS Chemical Neuroscience 8, 243–251 (2017).

18. Barandov, A., Bartelle, B. B., Williamson, C. G., Loucks, E. S., Lippard, S. J. & Jasanoff, A. Sensing intracellular calcium ions using a manganese-based MRI contrast agent. Nat Commun 10, 897 (2019).

19. Thiabaud, G. D., Schwalm, M., Sen, S., Barandov, A., Simon, J., Harvey, P., Spanoudaki, V., Müller, P., Sessler, J. L. & Jasanoff, A. Texaphyrin-Based Calcium Sensor for Multimodal Imaging. ACS Sens. (2023). doi:10.1021/acssensors.3c0138

20. Zhang, R., Li, L. S., Rao, B., Rong, H., Sun, M.-Y., Yao, J., Chen, R., Zhou, Q., Mennerick, S., Raman, B. & Wang, L. V. Multiscale photoacoustic tomography of neural activities with GCaMP calcium indicators. JBO 27, 096004 (2022).

21. Gottschalk, S., Degtyaruk, O., Mc Larney, B., Rebling, J., Hutter, M. A., Deán-Ben, X. L., Shoham, S. & Razansky, D. Rapid volumetric optoacoustic imaging of neural dynamics across the mouse brain. Nat Biomed Eng 3, 392–401 (2019).

22. Maresca, D., Lakshmanan, A., Abedi, M., Bar-Zion, A., Farhadi, A., Lu, G. J., Szablowski, J. O., Wu, D., Yoo, S. & Shapiro, M. G. Biomolecular Ultrasound and Sonogenetics. Annual Review of Chemical and Biomolecular Engineering 9, 229–252 (2018).

23. Rabut, C., Yoo, S., Hurt, R. C., Jin, Z., Li, H., Guo, H., Ling, B. & Shapiro, M. G. Ultrasound Technologies for Imaging and Modulating Neural Activity. Neuron 108, 93–110 (2020).

24. Shapiro, M. G., Goodwill, P. W., Neogy, A., Yin, M., Foster, F. S., Schaffer, D. V. & Conolly, S. M. Biogenic gas nanostructures as ultrasonic molecular reporters. Nature Nanotech 9, 311–316 (2014).

25. Bourdeau, R. W., Lee-Gosselin, A., Lakshmanan, A., Farhadi, A., Kumar, S. R., Nety, S. P. & Shapiro, M. G. Acoustic reporter genes for noninvasive imaging of microorganisms in mammalian hosts. Nature 553, 86–90 (2018).

26. Hurt, R. C., Buss, M. T., Duan, M., Wong, K., You, M. Y., Sawyer, D. P., Swift, M. B., Dutka, P., Barturen-Larrea, P., Mittelstein, D. R., Jin, Z., Abedi, M. H., Farhadi, A., Deshpande, R. & Shapiro, M. G. Genomically mined acoustic reporter genes for real-time in vivo monitoring of tumors and tumor-homing bacteria. Nat Biotechnol 1–13 (2023). doi:10.1038/s41587-022-01581-y

27. Farhadi, A., Ho, G. H., Sawyer, D. P., Bourdeau, R. W. & Shapiro, M. G. Ultrasound imaging of gene expression in mammalian cells. 7 (2019).

28. Lakshmanan, A., Farhadi, A., Nety, S. P., Lee-Gosselin, A., Bourdeau, R. W., Maresca, D. & Shapiro, M. G. Molecular Engineering of Acoustic Protein Nanostructures. ACS Nano 10, 7314–7322 (2016).

29. Maresca, D., Lakshmanan, A., Lee-Gosselin, A., Melis, J. M., Ni, Y.-L., Bourdeau, R. W., Kochmann, D. M. & Shapiro, M. G. Nonlinear ultrasound imaging of nanoscale acoustic biomolecules. Appl. Phys. Lett. 110, 073704 (2017).

30. Maresca, D., Sawyer, D. P., Renaud, G., Lee-Gosselin, A. & Shapiro, M. G. Nonlinear X-Wave Ultrasound Imaging of Acoustic Biomolecules. Phys. Rev. X 8, 041002 (2018).

31. Dutka, P., Metskas, L. A., Hurt, R. C., Salahshoor, H., Wang, T.-Y., Malounda, D., Lu, G. J., Chou, T.-F., Shapiro, M. G. & Jensen, G. J. Structure of Anabaena flos-aquae gas vesicles revealed by cryo-ET. Structure 31, 518–528.e6 (2023).

32. Huber, S. T., Terwiel, D., Evers, W. H., Maresca, D. & Jakobi, A. J. Cryo-EM structure of gas vesicles for buoyancy-controlled motility. Cell 186, 975–986.e13 (2023).

33. Lakshmanan, A., Jin, Z., Nety, S. P., Sawyer, D. P., Lee-Gosselin, A., Malounda, D., Swift, M. B., Maresca, D. & Shapiro, M. G. Acoustic biosensors for ultrasound imaging of enzyme activity. Nat Chem Biol 16, 988–996 (2020).

34. Heiles, B., Terwiel, D. & Maresca, D. The Advent of Biomolecular Ultrasound Imaging. Neuroscience 474, 122–133 (2021).

35. Nakai, J., Ohkura, M. & Imoto, K. A high signal-to-noise Ca2+ probe composed of a single green fluorescent protein. Nat Biotechnol 19, 137–141 (2001).

36. Clapperton, J. A., Martin, S. R., Smerdon, S. J., Gamblin, S. J. & Bayley, P. M. Structure of the Complex of Calmodulin with the Target Sequence of Calmodulin-Dependent Protein Kinase I: Studies of the Kinase Activation Mechanism. Biochemistry 41, 14669–14679 (2002).

37. Helassa, N., Zhang, X., Conte, I., Scaringi, J., Esposito, E., Bradley, J., Carter, T., Ogden, D., Morad, M. & Török, K. Fast-Response Calmodulin-Based Fluorescent Indicators Reveal Rapid Intracellular Calcium Dynamics. Sci Rep 5, 15978 (2015).

38. Lakshmanan, A., Lu, G. J., Farhadi, A., Nety, S. P., Kunth, M., Lee-Gosselin, A., Maresca, D., Bourdeau, R. W., Yin, M., Yan, J., Witte, C., Malounda, D., Foster, F. S., Schröder, L. & Shapiro, M. G. Preparation of biogenic gas vesicle nanostructures for use as contrast agents for ultrasound and MRI. Nat Protoc 12, 2050–2080 (2017).

39. Casey, J. R., Grinstein, S. & Orlowski, J. Sensors and regulators of intracellular pH. Nat Rev Mol Cell Biol 11, 50–61 (2010).

40. van der Linden, F. H., Mahlandt, E. K., Arts, J. J. G., Beumer, J., Puschhof, J., de Man, S. M. A., Chertkova, A. O., Ponsioen, B., Clevers, H., van Buul, J. D., Postma, M., Gadella, T. W. J. & Goedhart, J. A turquoise fluorescence lifetime-based biosensor for quantitative imaging of intracellular calcium. (Cell Biology, 2021). doi:10.1101/2021.06.21.449214

41. Gleichmann, M. & Mattson, M. P. Neuronal Calcium Homeostasis and Dysregulation. Antioxid Redox Signal 14, 1261–1273 (2011).

42. Huston, J. S., Levinson, D., Mudgett-Hunter, M., Tai, M. S., Novotný, J., Margolies, M. N., Ridge, R. J., Bruccoleri, R. E., Haber, E. & Crea, R. Protein engineering of antibody binding sites: recovery of specific activity in an anti-digoxin single-chain Fv analogue produced in Escherichia coli. Proceedings of the National Academy of Sciences 85, 5879–5883 (1988).

43. Inoue, M. Genetically encoded calcium indicators to probe complex brain circuit dynamics in vivo. Neuroscience Research 169, 2–8 (2021).

44. Park, H. Y., Kim, S. A., Korlach, J., Rhoades, E., Kwok, L. W., Zipfel, W. R., Waxham, M. N., Webb, W. W. & Pollack, L. Conformational changes of calmodulin upon Ca2+ binding studied with a microfluidic mixer. Proceedings of the National Academy of Sciences 105, 542–547 (2008).

45. Faas, G. C., Raghavachari, S., Lisman, J. E. & Mody, I. Calmodulin as a direct detector of Ca2+ signals. Nat Neurosci 14, 301–304 (2011).

46. Wang, Q., Zhang, P., Hoffman, L., Tripathi, S., Homouz, D., Liu, Y., Waxham, M. N. & Cheung, M. S. Protein recognition and selection through conformational and mutually induced fit. Proceedings of the National Academy of Sciences 110, 20545–20550 (2013).

47. Hellen, C. U. T. & Sarnow, P. Internal ribosome entry sites in eukaryotic mRNA molecules. Genes Dev. 15, 1593–1612 (2001).

48. Liu, C. & Hermann, T. E. Characterization of ionomycin as a calcium ionophore. Journal of Biological Chemistry 253, 5892–5894 (1978).

49. Ribeiro, C. M. P., McKay, R. R., Hosoki, E., Bird, G. St. J. & Putney, J. W. Effects of elevated cytoplasmic calcium and protein kinase C on endoplasmic reticulum structure and function in HEK293 cells. Cell Calcium 27, 175–185 (2000).

50. Dana, H., Mohar, B., Sun, Y., Narayan, S., Gordus, A., Hasseman, J. P., Tsegaye, G., Holt, G. T., Hu, A., Walpita, D., Patel, R., Macklin, J. J., Bargmann, C. I., Ahrens, M. B., Schreiter, E. R., Jayaraman, V., Looger, L. L., Svoboda, K. & Kim, D. S. Sensitive red protein calcium indicators for imaging neural activity. eLife 5, e12727 (2016).

51. Kofuji, P. & Araque, A. G-Protein-Coupled Receptors in Astrocyte–Neuron Communication. Neuroscience 456, 71–84 (2021).

52. Lämmermann, T. & Kastenmüller, W. Concepts of GPCR-controlled navigation in the immune system. Immunological Reviews 289, 205–231 (2019).

53. Capote, L. A., Mendez Perez, R. & Lymperopoulos, A. GPCR signaling and cardiac function. European Journal of Pharmacology 763, 143–148 (2015).

54. Mizuno, N. & Itoh, H. Functions and Regulatory Mechanisms of Gq-Signaling Pathways. Neurosignals 17, 42–54 (2009).

55. Sánchez-Fernández, G., Cabezudo, S., García-Hoz, C., Benincá, C., Aragay, A. M., Mayor, F. & Ribas, C. Gαq signalling: The new and the old. Cellular Signalling 26, 833–848 (2014).

56. Armbruster, B. N., Li, X., Pausch, M. H., Herlitze, S. & Roth, B. L. Evolving the lock to fit the key to create a family of G protein-coupled receptors potently activated by an inert ligand. Proceedings of the National Academy of Sciences 104, 5163–5168 (2007).

57. Nagai, Y., Miyakawa, N., Takuwa, H., Hori, Y., Oyama, K., Ji, B., Takahashi, M., Huang, X.-P., Slocum, S. T., DiBerto, J. F., Xiong, Y., Urushihata, T., Hirabayashi, T., Fujimoto, A., Mimura, K., English, J. G., Liu, J., Inoue, K., Kumata, J., Seki, C., Ono, M., Shimojo, M., Zhang, M.-R., Tomita, Y., Nakahara, J., Suhara, T., Takada, M., Higuchi, M., Jin, J., Roth, B. L. & Minamimoto, T. Deschloroclozapine, a potent and selective chemogenetic actuator enables rapid neuronal and behavioral modulations in mice and monkeys. Nat Neurosci 23, 1157–1167 (2020).

58. Nguyen, Q.-T., Schroeder, L. F., Mank, M., Muller, A., Taylor, P., Griesbeck, O. & Kleinfeld, D. An in vivo biosensor for neurotransmitter release and in situ receptor activity. Nat Neurosci 13, 127–132 (2010).

59. Muller, A., Joseph, V., Slesinger, P. A. & Kleinfeld, D. Cell-based reporters reveal in vivo dynamics of dopamine and norepinephrine release in murine cortex. Nat Methods 11, 1245–1252 (2014).

60. Xiong, H., Lacin, E., Ouyang, H., Naik, A., Xu, X., Xie, C., Youn, J., Wilson, B. A., Kumar, K., Kern, T., Aisenberg, E., Kircher, D., Li, X., Zasadzinski, J. A., Mateo, C., Kleinfeld, D., Hrabetova, S., Slesinger, P. A. & Qin, Z. Probing Neuropeptide Volume Transmission In Vivo by Simultaneous Near-Infrared Light-Triggered Release and Optical Sensing**. Angewandte Chemie 134, e202206122 (2022).

61. Doherty, J. E., Huye, L. E., Yusa, K., Zhou, L., Craig, N. L. & Wilson, M. H. Hyperactive piggyBac Gene Transfer in Human Cells and In Vivo. Hum Gene Ther 23, 311–320 (2012).

62. Loew, R., Heinz, N., Hampf, M., Bujard, H. & Gossen, M. Improved Tet-responsive promoters with minimized background expression. BMC Biotechnol 10, 81 (2010).

63. Rahal, L., Thibaut, M., Rivals, I., Claron, J., Lenkei, Z., Sitt, J. D., Tanter, M. & Pezet, S. Ultrafast ultrasound imaging pattern analysis reveals distinctive dynamic brain states and potent sub-network alterations in arthritic animals. Sci Rep 10, 10485 (2020).

64. Brunner, C., Grillet, M., Urban, A., Roska, B., Montaldo, G. & Macé, E. Whole-brain functional ultrasound imaging in awake head-fixed mice. Nat Protoc 16, 3547–3571 (2021).

65. Roth, B. L. DREADDs for Neuroscientists. Neuron 89, 683–694 (2016).

66. Mace, E., Montaldo, G., Cohen, I., Baulac, M., Fink, M. & Tanter, M. Functional ultrasound imaging of the brain. Nat Methods 8, 662–4 (2011).

67. Rabut, C., Correia, M., Finel, V., Pezet, S., Pernot, M., Deffieux, T. & Tanter, M. 4D functional ultrasound imaging of whole-brain activity in rodents. Nat Methods 16, 994–997 (2019).

68. Duan, M., Dev, I., Lu, A., You, M. Y. & Shapiro, M. G. Stoichiometric expression of messenger polycistrons by eukaryotic ribosomes (SEMPER) for compact, ratio-tunable multi-gene expression from single mRNAs. (Synthetic Biology, 2023). doi:10.1101/2023.05.26.541240

69. Wang, C., Chen, X., Wang, L., Makihata, M., Liu, H.-C., Zhou, T. & Zhao, X. Bioadhesive ultrasound for long-term continuous imaging of diverse organs. Science 377, 517–523 (2022).

70. Hu, H., Huang, H., Li, M., Gao, X., Yin, L., Qi, R., Wu, R. S., Chen, X., Ma, Y., Shi, K., Li, C., Maus, T. M., Huang, B., Lu, C., Lin, M., Zhou, S., Lou, Z., Gu, Y., Chen, Y., Lei, Y., Wang, X., Wang, R., Yue, W., Yang, X., Bian, Y., Mu, J., Park, G., Xiang, S., Cai, S., Corey, P. W., Wang, J. & Xu, S. A wearable cardiac ultrasound imager. Nature 613, 667–675 (2023).

71. Errico, C., Pierre, J., Pezet, S., Desailly, Y., Lenkei, Z., Couture, O. & Tanter, M. Ultrafast ultrasound localization microscopy for deep super-resolution vascular imaging. Nature 527, 499–502 (2015).

72. Sofroniew, N. J., Flickinger, D., King, J. & Svoboda, K. A large field of view two-photon mesoscope with subcellular resolution for in vivo imaging. eLife 5, e14472 (2016).

73. Ahrens, M. B., Orger, M. B., Robson, D. N., Li, J. M. & Keller, P. J. Whole-brain functional imaging at cellular resolution using light-sheet microscopy. Nat Methods 10, 413–420 (2013).

74. Skocek, O., Nöbauer, T., Weilguny, L., Martínez Traub, F., Xia, C. N., Molodtsov, M. I., Grama, A., Yamagata, M., Aharoni, D., Cox, D. D., Golshani, P. & Vaziri, A. High-speed volumetric imaging of neuronal activity in freely moving rodents. Nat Methods 15, 429–432 (2018).

75. Gibson, D. G., Young, L., Chuang, R.-Y., Venter, J. C., Hutchison, C. A. & Smith, H. O. Enzymatic assembly of DNA molecules up to several hundred kilobases. Nat Methods 6, 343–345 (2009).

76. Salahshoor, H., Yao, Y., Dutka, P., Nyström, N. N., Jin, Z., Min, E., Malounda, D., Jensen, G. J., Ortiz, M. & Shapiro, M. G. Geometric effects in gas vesicle buckling under ultrasound. Biophysical Journal 121, 4221–4228 (2022).

77. Mastronarde, D. N. Automated electron microscope tomography using robust prediction of specimen movements. Journal of Structural Biology 152, 36–51 (2005).

78. Punjani, A., Rubinstein, J. L., Fleet, D. J. & Brubaker, M. A. cryoSPARC: algorithms for rapid unsupervised cryo-EM structure determination. Nat Methods 14, 290–296 (2017).

79. Mirdita, M., Schütze, K., Moriwaki, Y., Heo, L., Ovchinnikov, S. & Steinegger, M. ColabFold: making protein folding accessible to all. Nat Methods 19, 679–682 (2022).

80. Goddard, T. D., Huang, C. C., Meng, E. C., Pettersen, E. F., Couch, G. S., Morris, J. H. & Ferrin, T. E. UCSF ChimeraX: Meeting modern challenges in visualization and analysis. Protein Science 27, 14–25 (2018).

81. Maresca, D., Payen, T., Lee-Gosselin, A., Ling, B., Malounda, D., Demené, C., Tanter, M. & Shapiro, M. G. Acoustic biomolecules enhance hemodynamic functional ultrasound imaging of neural activity. NeuroImage 209, 116467 (2020).

82. Ling, B., Lee, J., Maresca, D., Lee-Gosselin, A., Malounda, D., Swift, M. B. & Shapiro, M. G. Biomolecular Ultrasound Imaging of Phagolysosomal Function. ACS Nano 14, 12210–12221 (2020).

83. Demené, C., Deffieux, T., Pernot, M., Osmanski, B.-F., Biran, V., Gennisson, J.-L., Sieu, L.-A., Bergel, A., Franqui, S., Correas, J.-M., Cohen, I., Baud, O. & Tanter, M. Spatiotemporal Clutter Filtering of Ultrafast Ultrasound Data Highly Increases Doppler and fUltrasound Sensitivity. IEEE Transactions on Medical Imaging 34, 2271–2285 (2015).

84. Mace, E., Montaldo, G., Osmanski, B.-F., Cohen, I., Fink, M. & Tanter, M. Functional ultrasound imaging of the brain: theory and basic principles. IEEE Transactions on Ultrasonics, Ferroelectrics, and Frequency Control 60, 492–506 (2013).

