## Supplementary Materials for "Ultrasonic reporters of calcium for deep tissue imaging of cellular signals"

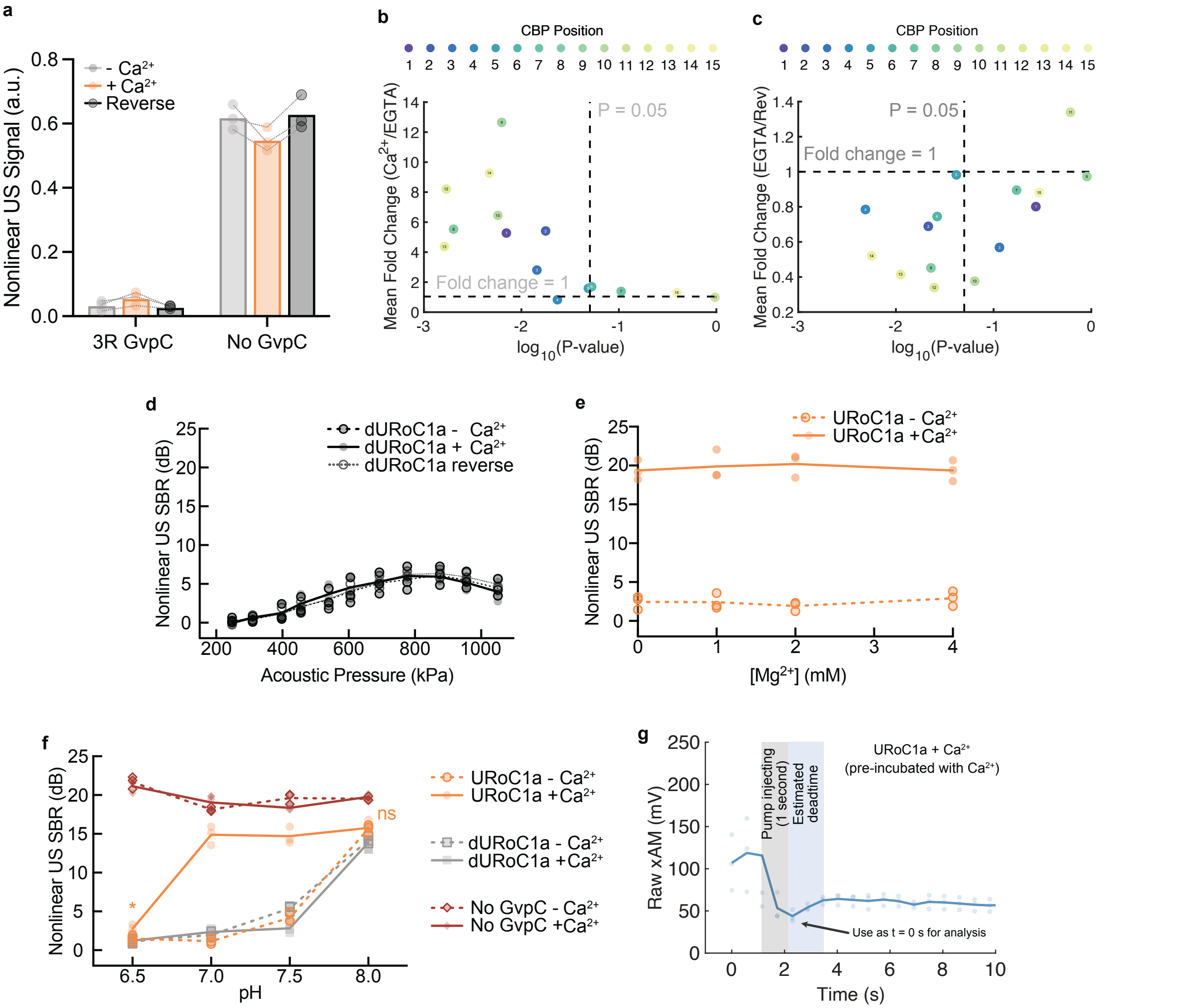


**Supplementary Figure 1. URoC screening and URoC1a characterization**. (**a**) Nonlinear signal after background subtraction and GV concentration normalization of GVs with 3-repeat Ana GvpC (3R GvpC) or no GvpC after incubation with EGTA, calcium or first with calcium and then with EGTA. (**b**, **c**) The nonlinear signal mean fold change of URoC variants with different CBP insertion sites as a function the p-value, comparing conditions with calcium and with EGTA (**b**) or first with calcium and then reversed with EGTA and only with EGTA (**c**). The vertical dash lines represent the significance threshold of $p = 0.05$ and the horizontal lines represent no calcium-dependent change (**b**) or fully reversible (**c**). (**d**) Nonlinear SBR in dB scale as a function of applied acoustic pressure for control dURoC1a GVs (with all EF hands knocked out) after incubation with 200 μM $CaCl_2$, 5 mM EGTA or first with 200 μM $CaCl_2$ and then with 5 mM EGTA. Solid curves represent the mean of all biological replicates. (**e**) Nonlinear SBR in dB scale as a function of magnesium concentration for URoC1a after incubation with 1 mM $CaCl_2$ (solid lines) or 5 mM EGTA (dash lines). Curves represent the mean of all biological replicates. (**f**) Nonlinear SBR in dB scale as a function of pH for URoC1a, control GVs or GVs without GvpC after incubation with 200 µM $CaCl_2$ (solid lines) or 5 mM EGTA (dash lines). (**g**) Nonlinear ultrasound signal of URoC1a preincubated with 200 µM $CaCl_2$ as a function of time after being mixed into 200 µM $CaCl_2$. The time needed for the signal to become stable was used to estimate the dead time for the kinetics measurement. Ultrasound data were acquired with xAM at 547 kPa for (**a-c**, **e-f**) and at 472 kPa for (**g**). Curves represent the mean of all biological replicates. N = 3 biological replicates for all panels and each biological replicate has 2 technical replicates for **d, e, f**. Dots represent individual measurement of each replicate for **a** and **g**, or the mean of two technical replicates for **d, e, f**.

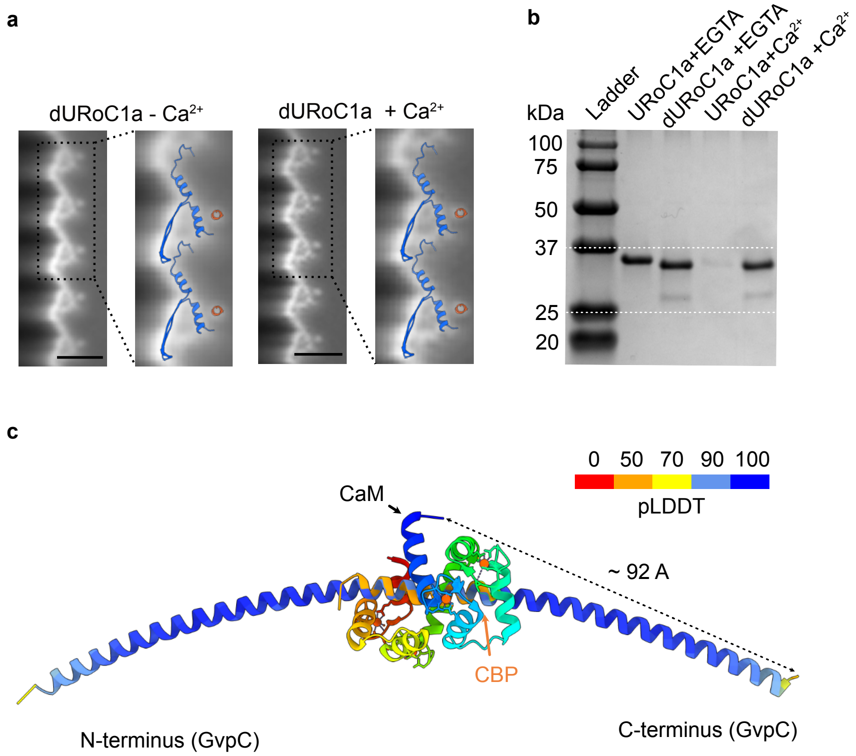


**Supplementary Figure 2. Molecular mechanism of URoC.** (**a**) Cryo-EM 2D density map of the side view of the dURoC1a GV shell incubated with 200 µM $CaCl_2$ or 5 mM EGTA prior to freezing and an integrative model of the Ana GvpA:GvpC (PDB: 8GBS)[31] complex was overlaid on the GV shell density in the 2D class averages. (**b**) Coomassie-stained SDS-PAGE gel of $OD_{500nm}$-matched URoC1a GVs and dURoC1a with unbound GvpC molecules removed through buoyancy purification after incubation with calcium or EGTA at 37°C. (**c**) Structure prediction of calcium-saturated CaM bound to the URoC GvpC without the linker. The GvpC was colored by the pLDDT confidence from AlphaFold prediction. The CaM was colored in rainbow from blue to red (N-terminus to C-terminus) and the CBP in the CaM-CBP complex was colored orange. The dash line indicates the distance between the center carbon of the C-terminal glycine of GvpC and the that of the N-terminal aspartic acid of CaM.

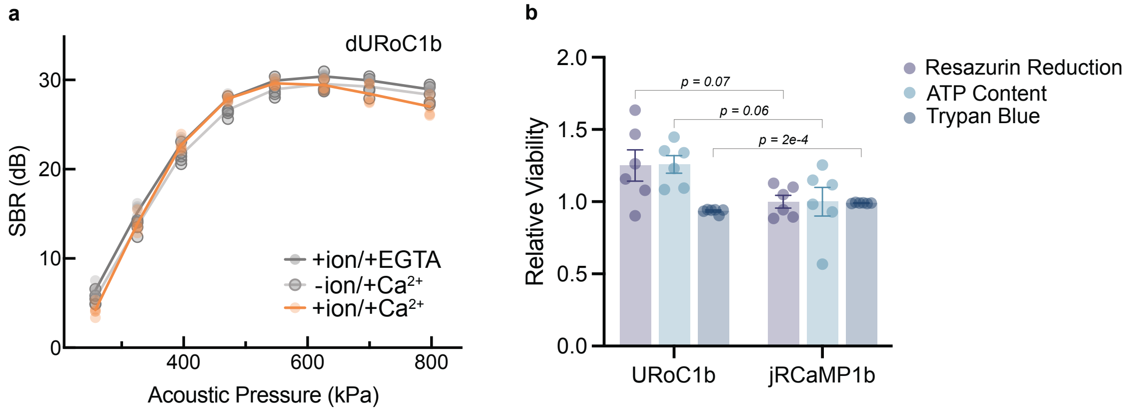


**Supplementary Figure 3. Ultrasound imaging of control non-calcium-sensing GVs in mammalian cells and viability assay.** (**a**) Nonlinear SBR in dB scale as a function of applied acoustic pressure for cells expressing control dURoC1b after incubation with ionomycin and EGTA, ionomycin and calcium or calcium only. (**b**) Viability assays of cells transiently expressing URoC1b or jRCaMP1b. The data were normalized to the mean of each measurement done with jRCaMp1b. The difference in viability measured by Trypan Blue is statistically different but both are above 93% (93.5% for URoC1b and 99% for jRCaMP1b).

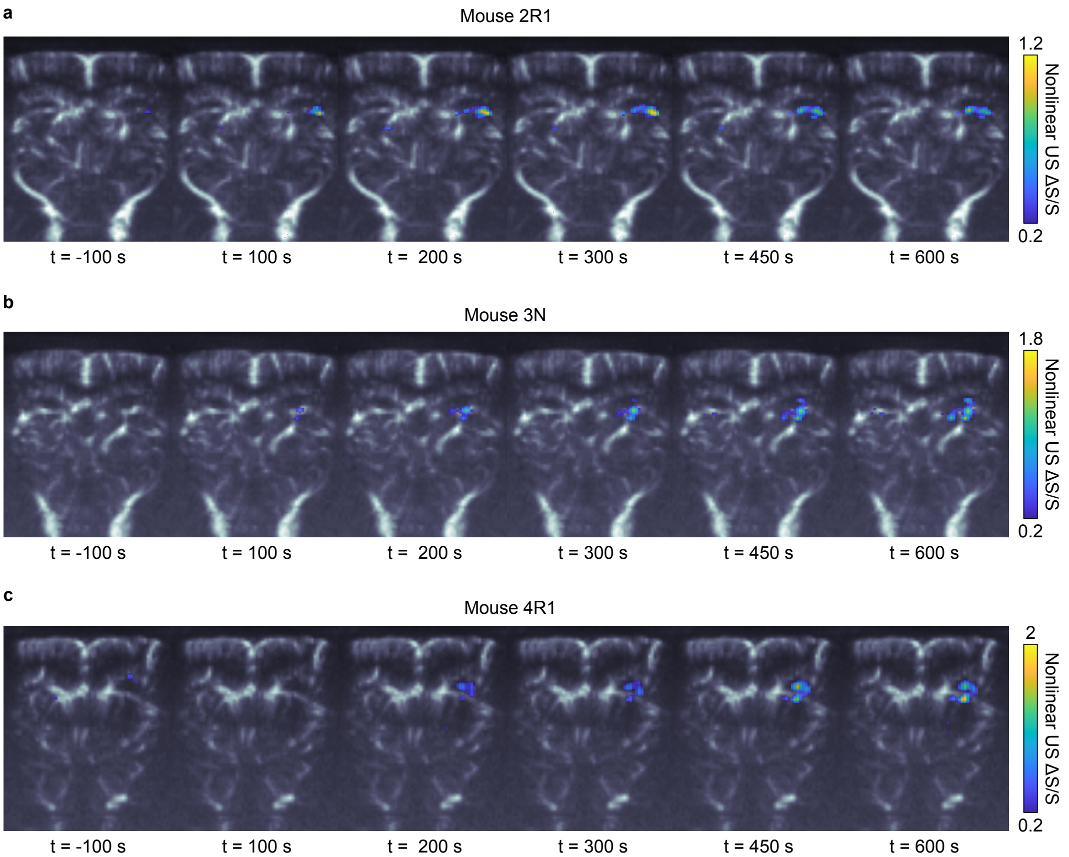


**Supplementary Figure 4. Ultrasound imaging of ligand-induced GPCR-driven calcium dynamics *in vivo*.** (**a-c**) Representative nonlinear US images of DRUM and $DRUM_{mut}$ implants in the brain of mouse 2R1 (**a**), 3N (**b**) and 4R1 (**c**). The relative signal change at t = -100/100/200/300/450/600 s compared to t = 0 s was overlaid on a Doppler image. The Scale bar = 1 mm. Color bars represent ultrasound signal intensity in the dB scale. The intensity of the Doppler image represents the blood volume that circulated in each voxel within an integration time of 400 ms (see Methods). Imaging was conducted through polymer cranial windows.

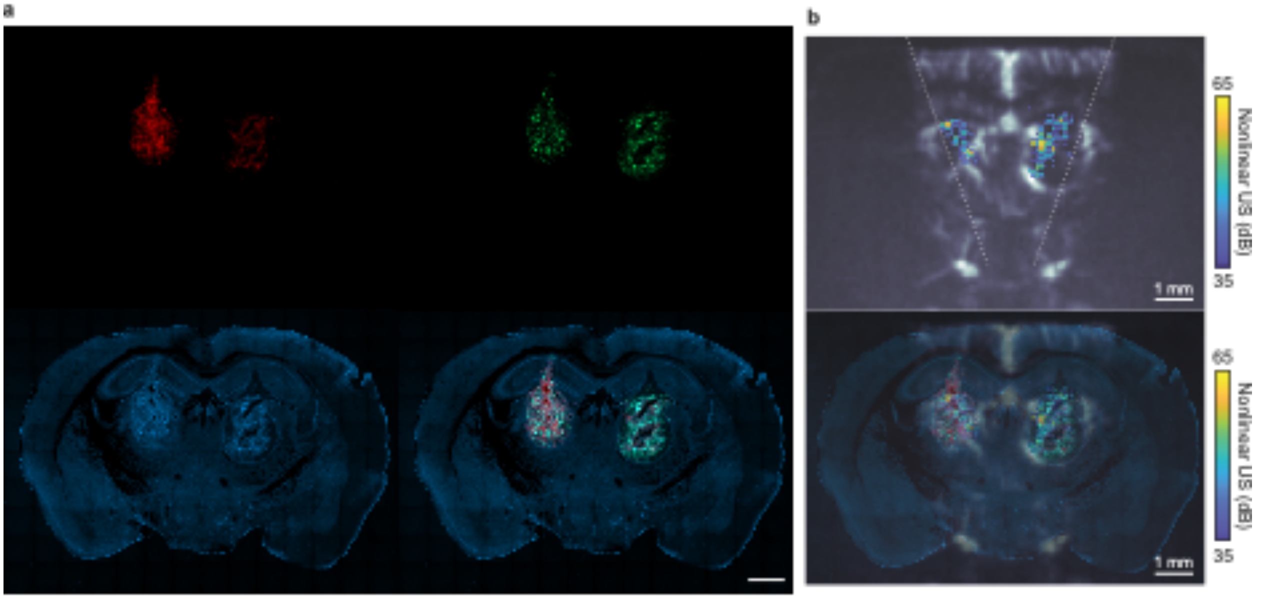


**Supplementary Figure 5. Immunofluorescence characterization of DRUM and $DRUM_{mut}$ implants *in vivo*.** (**a**) Representative immunofluorescence micrograph of a 100-µm-thin brain section. Red color shows mCherry fluorescence from the hM3D(Gq) receptor (direct fusion); green color shows GFP fluorescence from the *gvpNV* genes (P2A chained); blue shows DAPI nuclear stain. (**b**) Top: the nonlinear ultrasound image of the baseline signals from the implants overlaid on a Doppler image for the same mouse before DCZ injection (t = 0 s). The dashed line indicates the effective field of view of xAM due to the side aperture partially blocked by the edge of the skull/cranial window. Bottom: The ultrasound image overlaid with the fluorescence micrograph. All the scale bars = 1 mm. Color bars represent ultrasound signal intensity in the dB scale. The nonlinear ultrasound signal was acquired with xAM at 426 kPa (calibrated in water) and through polymer cranial windows.

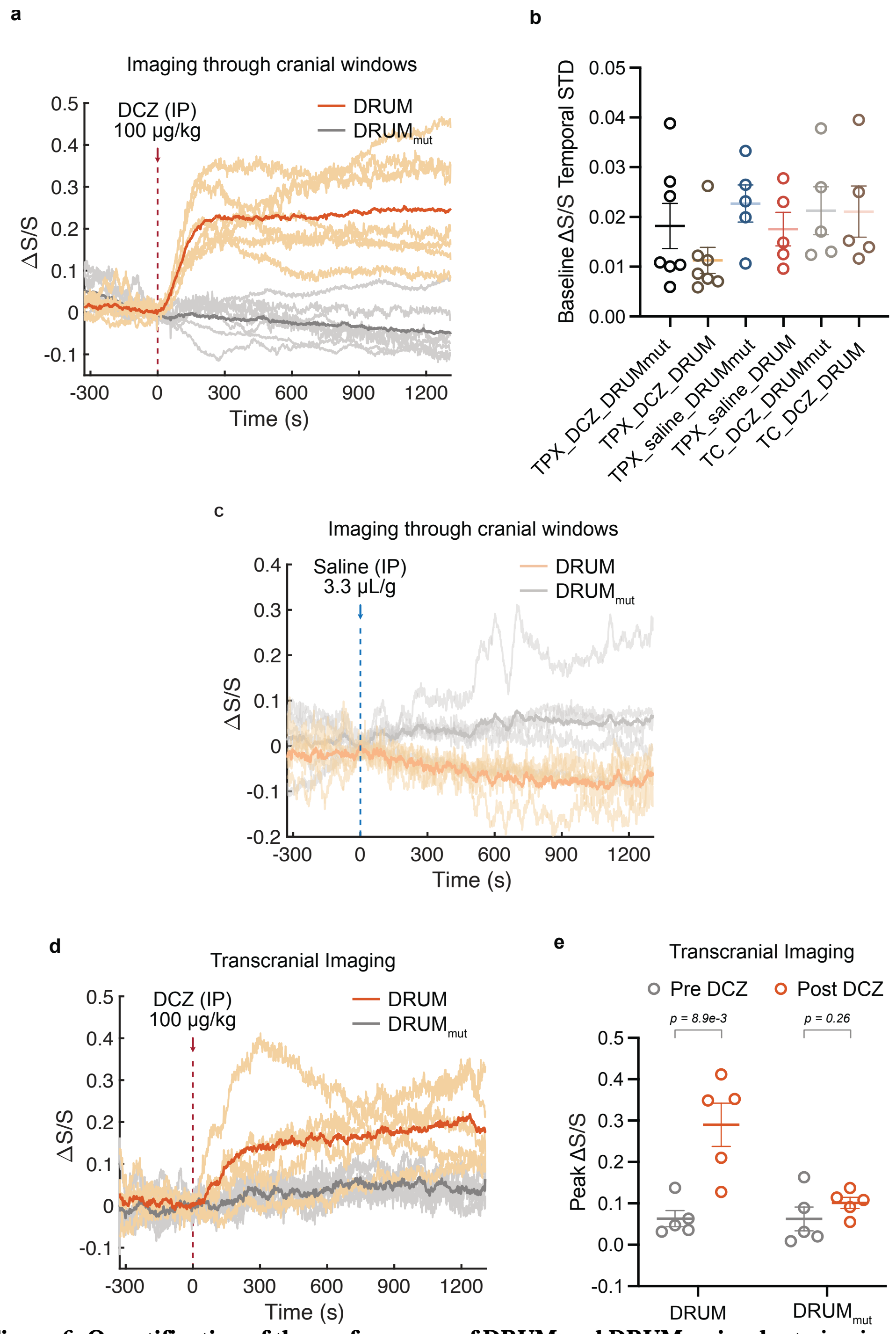


**Supplementary Figure 6. Quantification of the performance of DRUM and DRUM$_{mut}$ implants *in vivo*.**
(**a**) Time traces of nonlinear ultrasound contrast of the implanted cells through the cranial windows. DCZ was injected I.P. at t = 0 s at the dash line. Dark curve represents the mean of all traces and light curves represent data from individual animals. (**b**) Temporal standard deviation of the baseline relative change of nonlinear signal of DRUM or DRUM$_{mut}$ implants from t = -327 s to t = 0 s. N = 7 for imaging through TPX® cranial window and stimulated with DCZ, N = 5 for the experiment with TPX® cranial window and saline stimulation, and transcranially (TC) with DCZ stimulation. Dots = individual trial, lines represent mean and error bars represent SEM. (**c**) Time traces of nonlinear ultrasound contrast of the implanted cells through the cranial windows. Saline was injected I.P. at t = 0 s at the dash line. Dark curve represents the mean of all traces and light curves represent data from individual animals. (**d**) Time traces of nonlinear ultrasound contrast of the implanted cells through the intact skulls. DCZ was injected I.P. at t = 0 s at the dash line. (**e**) Peak relative signal change of DRUM and DRUM$_{mut}$ implants before and after DCZ stimulation when imaging through the intact skulls. Dark curve represents the mean of all traces and light curves represent data from individual animals. Lines = mean and error bars = SEM. N = 7 mice for (**a**) and N = 5 mice for (**c-e**).

| Plasmid Name | Purpose | Reference Information |
|---|---|---|
| jbC-76_pET28a_GvpC-r45d | Bacterial expression of wild type 3-repeat Ana GvpC with 6xHis tag on the C-terminus for purification in pET28a backbone | Fig. 1d; Fig. S1a (available on Addgene upon publication). |
| jbC-78_pET-28a_URoC0-p1 | Bacterial expression of URoC variant with 8xG4S, CaM-EF1KO and CaMKI CBP substitute-insertion starting from the #1 residue of the second repeat of a 3-repeat GvpC in pET28a backbone | Fig. 1d; Fig. S1b-c. |
| jbC-79_pET-28a_URoC0-p2 | Bacterial expression of URoC variant with 8xG4S, CaM-EF1KO and CaMKI CBP substitute-insertion starting from the #2 residue of the second repeat of a 3-repeat GvpC in pET28a backbone | Fig. 1d; Fig. S1b-c |
| jbC-80_pET-28a_URoC0-p3 | Bacterial expression of URoC variant with 8xG4S, CaM-EF1KO and CaMKI CBP substitute-insertion starting from the #3 residue of the second repeat of a 3-repeat GvpC in pET28a backbone | Fig. 1d; Fig. S1b-c. |
| jbC-81_pET-28a_URoC0-p4 | Bacterial expression of URoC variant with 8xG4S, CaM-EF1KO and CaMKI CBP substitute-insertion starting from the #4 residue of the second repeat of a 3-repeat GvpC in pET28a backbone | Fig. 1d; Fig. S1b-c. |
| jbC-8_pET-28a_URoC0-p5 | Bacterial expression of URoC variant with 8xG4S, CaM-EF1KO and CaMKI CBP substitute-insertion starting from the #5 residue of the second repeat of a 3-repeat GvpC in pET28a backbone | Fig. 1d; Fig. S1b-c. |
| jbC-82_pET-28a_URoC0-p6 | Bacterial expression of URoC variant with 8xG4S, CaM-EF1KO and CaMKI CBP substitute-insertion starting from the #6 residue of the second repeat of a 3-repeat GvpC in pET28a backbone | Fig. 1d; Fig. S1b-c. |
| jbC-7_pET-28a_URoC0-p7 | Bacterial expression of URoC variant with 8xG4S, CaM-EF1KO and CaMKI CBP substitute-insertion starting from the #7 residue of the second repeat of a 3-repeat GvpC in pET28a backbone | Fig. 1d; Fig. S1b-c. |
| jbC-83_pET-28a_URoC0-p8 | Bacterial expression of URoC variant with 8xG4S, CaM-EF1KO and CaMKI CBP substitute-insertion starting from the #8 residue of the second repeat of a 3-repeat GvpC in pET28a backbone | Fig. 1d; Fig. S1b-c. |
| jbC-6_pET-28a_URoC1a | Bacterial expression of URoC1a with 8xG4S, CaM-EF1KO and CaMKI CBP substitute-insertion starting from the #9 residue of the second repeat of a 3-repeat GvpC in pET28a backbone | Fig. 1d-g, j; Fig. 2a-g; Fig. S1b-c, e-g; Fig. S2b (available on Addgene upon publication). |
| jbC-84_pET-28a_URoC0-p10 | Bacterial expression of URoC variant with 8xG4S, CaM-EF1KO and CaMKI CBP substitute-insertion starting from the #10 residue of the second repeat of a 3-repeat GvpC in pET28a backbone | Fig. 1d; Fig. S1b-c. |
| jbC-85_pET-28a_URoC0-p11 | Bacterial expression of URoC variant with 8xG4S, CaM-EF1KO and CaMKI CBP substitute-insertion starting from the #11 residue of the second repeat of a 3-repeat GvpC in pET28a backbone | Fig. 1d; Fig. S1b-c. |
| jbC-86_pET-28a_URoC0-p12 | Bacterial expression of URoC variant with 8xG4S, CaM-EF1KO and CaMKI CBP substitute-insertion starting from the #12 residue of the second repeat of a 3-repeat GvpC in pET28a backbone | Fig. 1d; Fig. S1b-c. |
| jbC-87_pET-28a_URoC0-p13 | Bacterial expression of URoC variant with 8xG4S, CaM-EF1KO and CaMKI CBP substitute-insertion starting from the #13 residue of the second repeat of a 3-repeat GvpC in pET28a backbone | Fig. 1d; Fig. S1b-c. |
| jbC-88_pET-28a_URoC0-p14 | Bacterial expression of URoC variant with 8xG4S, CaM-EF1KO and CaMKI CBP substitute-insertion starting from the #14 residue of the | Fig. 1d; Fig. S1b-c. |

| | | |
|---|---|---|
| | second repeat of a 3-repeat GvpC in pET28a backbone | |
| jbC-89_pET-28a_URoC0-p15 | Bacterial expression of URoC variant with 8xG4S, CaM-EF1KO and CaMKI CBP substitute-insertion starting from the #15 residue of the second repeat of a 3-repeat GvpC in pET28a backbone | Fig. 1d; Fig. S1b-c. |
| jbC-3_pET-28a_dURoC1a | Bacterial expression of control URoC1 with 8xG4S, CaM with all EF hands mutated out and CaMKI CBP substitute-insertion at the #9 residue of the second repeat of a 3-repeat GvpC in pET28a backbone | Fig. S1d, f; Fig. S2a-b (available on Addgene upon publication). |
| jbC-41_pET-28a_URoC1-FL2 | Bacterial expression of URoC1 variant with 2xG4S, CaM-EF1KO and CaMKI CBP substitute-insertion starting from the #9 residue of the second repeat of a 3-repeat GvpC in pET28a backbone | Fig. 2d-e. |
| jbC-38_pET-28a_URoC1-FL4 | Bacterial expression of URoC1 variant with 4xG4S, CaM-EF1KO and CaMKI CBP substitute-insertion starting from the #9 residue of the second repeat of a 3-repeat GvpC in pET28a backbone | Fig. 2d-e. |
| jbC-47_pET-28a_URoC1-FL12 | Bacterial expression of URoC1 variant with 12xG4S, CaM-EF1KO and CaMKI CBP substitute-insertion starting from the #9 residue of the second repeat of a 3-repeat GvpC in pET28a backbone | Fig. 2d-e. |
| jbC-49_pET-28a_URoC1-FL16 | Bacterial expression of URoC1 variant with 16xG4S, CaM-EF1KO and CaMKI CBP substitute-insertion starting from the #9 residue of the second repeat of a 3-repeat GvpC in pET28a backbone | Fig. 2d-e. |
| jbC-50_pET-28a_URoC1-EF2KO | Bacterial expression of URoC1 variant with 8xG4S, CaM-EF2KO and CaMKI CBP substitute-insertion starting from the #9 residue of the second repeat of a 3-repeat GvpC in pET28a backbone | Fig. 2f-g. |
| jbC-51_pET-28a_URoC1-EF3KO | Bacterial expression of URoC1 variant with 8xG4S, CaM-EF3KO and CaMKI CBP substitute-insertion starting from the #9 residue of the second repeat of a 3-repeat GvpC in pET28a backbone | Fig. 2f-g. |
| jbC-52_pET-28a_URoC1-EF4KO | Bacterial expression of URoC1 variant with 8xG4S, CaM-EF4KO and CaMKI CBP substitute-insertion starting from the #9 residue of the second repeat of a 3-repeat GvpC in pET28a backbone | Fig. 2f-g. |
| jbC-43_pET-28a_URoC1-nKO | Bacterial expression of URoC1 variant with 8xG4S, CaM6f and CaMKI CBP substitute-insertion starting from the #9 residue of the second repeat of a 3-repeat GvpC in pET28a backbone | Fig. 2f-g. |
| jbC-53_pET-28a_URoC1b | Bacterial expression of URoC1b with 2xG4S, CaM-EF2KO and CaMKI CBP substitute-insertion starting from the #9 residue of the second repeat of a 3-repeat GvpC in pET28a backbone | Fig. 2h-i (available on Addgene upon publication). |
| jmT-89_pCMV-A-IRES-URoC1b-WPRE-hGH | Transient mammalian expression of gvpA gene and URoC1b gvpC | Fig. 3c-e; Fig. S3b (available on Addgene upon publication). |
| jmT-80_pCMV-A-IRES-dURoC1b-WPRE-hGH | Transient mammalian expression of gvpA gene and dURoC1b gvpC | Fig. 3e, Fig.S3a (available on Addgene upon publication). |
| jmL-95_pLV-EF1a-hM3D(Gq)-mCherry-IRES-PuroR-WPRE | Lenti viral construct encoding DREADD receptor hM3D(Gq) tethered with mCherry and puromycin resistance marker. | Fig. 4d-g; Fig. S4-6 (available on Addgene upon publication). |
| jmT-88_pPB-TRE-A-IRES-URoC1b-WPRE-mEF1a-rtTA-T2A-HygR | PiggyBac transposon plasmid for inducible expression of gvpA and URoC1b gvpC genes with a hygromycin resistance gene | Fig. 4d-g; Fig. S4-6 (available on Addgene upon publication). |

| | | |
|---|---|---|
| jmT-87_pPB-TRE-A-IRES-dURoC1b-WPRE-mEF1a-rtTA-T2A-HygR | PiggyBac transposon plasmid for inducible expression of gvpA and dURoC1b gvpC genes with a hygromycin resistance gene | Fig. 4d-g; Fig. S4-6 (available on Addgene upon publication). |
| jmT-45_pPB-TRE-NV(allP2A)-emGFP-WPRE-mEF1a-BSD | PiggyBac transposon plasmid for inducible expression of Ana GV chaperone genes with a blasticidin resistance gene | Fig. 4d-g; Fig. S4-6 (available on Addgene upon publication). |
| mARGAna (cassette 2, transient) | Second-generation mammalian acoustic reporter gene (cassette 2), used for transient transfection in this work. | Addgene #197589 |
| mARGAna (cassette 1, transient) | Second-generation mammalian acoustic reporter gene (cassette 1), used as cloning backbone. | Addgene #197588 |
| mARGAna (cassette 1) | Second-generation mammalian acoustic reporter gene (cassette 1) on the PiggyBac transposon plasmid, used as cloning backbones. | Addgene #191341 |
| mARGAna (cassette 2) | Second-generation mammalian acoustic reporter gene (cassette 2) on the PiggyBac transposon plasmid, used as cloning backbones. | Addgene #191342 |
| pAAV-hSyn-hM3D(Gq)-mCherry | Gq-coupled hM3D DREADD fused with mCherry under the control of human synapsin promoter, used as cloning templates to generate jmL-95. | Addgene #50474 |

**Supplementary Table 1: List and features of genetic constructs used in this study.**
pAAV-hSyn-hM3D(Gq)-mCherry[56] was a gift from Bryan Roth (Addgene plasmid # 50474 ; http://n2t.net/addgene:50474 ; RRID:Addgene_50474)
